# Mapping the topography of spatial gene expression with interpretable deep learning

**DOI:** 10.1101/2023.10.10.561757

**Authors:** Uthsav Chitra, Brian J. Arnold, Hirak Sarkar, Cong Ma, Sereno Lopez-Darwin, Kohei Sanno, Benjamin J. Raphael

## Abstract

Spatially resolved transcriptomics technologies provide high-throughput measurements of gene expression in a tissue slice, but the sparsity of this data complicates the analysis of spatial gene expression patterns such as gene expression gradients. We address these issues by deriving a *topographic map* of a tissue slice—analogous to a map of elevation in a landscape—using a novel quantity called the *isodepth*. Contours of constant isodepth enclose spatial domains with distinct cell type composition, while gradients of the isodepth indicate spatial directions of maximum change in gene expression. We develop GASTON, an unsupervised and interpretable deep learning algorithm that simultaneously learns the isodepth, spatial gene expression gradients, and piecewise linear functions of the isodepth that model both continuous gradients and discontinuous spatial variation in the expression of individual genes. We validate GASTON by showing that it accurately identifies spatial domains and marker genes across several biological systems. In SRT data from the brain, GASTON reveals gradients of neuronal differentiation and firing, and in SRT data from a tumor sample, GASTON infers gradients of metabolic activity and epithelial-mesenchymal transition (EMT)-related gene expression in the tumor microenvironment.

## 1 Introduction

Gene expression varies substantially across a tissue, due to both the spatial organization of cell types within a tissue and localized changes in cell state through processes such as development, differentiation, and intercellular communication [160]. Many genes display sharp, discontinuous changes in expression in certain areas of a tissue, often near the boundaries of distinct *spatial domains* containing different combinations of cell types. For example, different cortical and neocortical layers of the brain are distinguished by the presence and absence of expression of certain marker genes [124, 96]. Gene expression may also vary *continuously* in a tissue, forming gene expression *“gradients”* that distinguish different cell types or states and drive fundamental biological processes including development [6, 55, 48, 117] and cellular communication [148, 138]. For instance, gene expression gradients underlie the functional heterogeneity of neurons in the hippocampus [160, 21] and hepatocytes in individual liver lobules [9, 25]. In tumors, gene expression may vary continuously with the distance to the surrounding stroma due to oxygen gradients or cellular interactions [125, 12].

Spatially resolved transcriptomics (SRT) technologies produce high-throughput measurements of spatial gene expression, quantifying the number of RNA transcripts at thousands in a tissue slice [93, 88, 101, 111, 134, 139]. These SRT technologies enable the inference of spatial domains in tissues as well as the identification of genes and cell types with continuous and discontinuous spatial patterns of expression within and across spatial domains. However, SRT technologies typically yield sparse measurements of the transcriptome: current whole-transcriptome sequencing-based technologies [1, 116, 127, 22, 78] have limited coverage (≈500-5,000 unique molecular identifiers (UMIs) per location) while imaging-based technologies measure a much smaller and targeted panel of transcripts (typically 100-1,000 transcripts) [61, 143, 162, 91, 52]. This sparsity markedly complicates the analysis of spatial gene expression.

Numerous computational approaches have been developed to identify spatial domains and/or genes with spatially varying expression from SRT data. These methods typically leverage correlations between expression measurements at nearby spatial locations to overcome the sparse measurements at individual locations. Many methods focus on the identification of distinct spatial domains by partitioning tissues into subregions having large, discontinuous changes in gene expression, e.g. [168, 58, 32, 104, 81, 153, 76, 167, 53], but do not model continuous gene expression gradients within these regions. Several other methods instead test whether the expression of an individual gene varies spatially by fitting a function to the observed transcript counts at spatial locations [132, 130, 171, 18, 145]. However, these methods cannot distinguish continuous gradients within spatial domains from discontinuous changes in expression between domains. More generally, neither approach models the geometry of a tissue slice using a coordinate system that describes both the boundaries of spatial domains and the *relative* position of spatial locations within these domains, thus greatly limiting their ability to identify continuous gradients of gene expression.

We introduce *gene expression topography*, a fundamentally different approach to modeling spatial variation in gene expression. We derive a *“topographic map”* of a tissue slice using the *isodepth*, a 1-dimensional coordinate over the tissue slice which describes both the arrangement of spatial domains and the relative position of each spatial location within its corresponding spatial domain. Thus, just as the topographic map of a landscape demarcates mountains and valleys by their elevation, our topographic map of gene expression delineates spatial domains by their isodepth. Moreover, like the elevation of a landscape, the isodepth varies continuously over a tissue slice, providing a coordinate to model continuous variation in the expression of individual genes. In particular, our topographic map describes *gene expression gradients*, similar to how a topographic map of elevation shows whether a direction is a steep ascent or a flat plateau.

We develop Gradient Analysis of Spatial Transcriptomics Organization with Neural networks (GASTON), an unsupervised and interpretable deep neural network algorithm that learns the isodepth of a tissue slice, the vector field of spatial gradients of gene expression, and spatial expression functions for individual genes directly from SRT data. In particular, GASTON models gene expression as a *piecewise linear* function of the isodepth, thus describing both continuous gradients and sharp discontinuities in gene expression. We demonstrate that the isodepth and spatial gradients learned by GASTON reveal the geometry and continuous gene expression gradients of multiple tissues across multiple SRT technologies including 10x Genomics Visium [1], Slide-SeqV2 [116, 127], and Stereo-Seq [22]. On SRT data from the mouse and human brain, we show that GASTON more accurately identifies spatial domains and marker genes compared to existing methods, derives maps of spatial variation in cell type organization, and uncovers spatial gradients of neuronal firing and differentiation. Using SRT data from a colorectal tumor sample, we demonstrate that GASTON identifies gradients of metabolic activity in the tumor interior, and gradients of epithelialmesenchymal transition (EMT)-related gene expression at the tumor-stroma boundary.

## 2 Results

### 2.1 GASTON learns the topography of a tissue slice using interpretable deep learning

We introduce the *isodepth d*, a scalar quantity that models the *“topography”* of a tissue slice and is analogous to the elevation in a topographic map of a land surface. A small number of contours of equal isodepth *d* partition the tissue slice into spatial domains, while the intermediate isodepth contours define the relative position of a location within a domain. Moreover, the gradient ∇*d* of the isodepth *d* at each location describes the *spatial gradient*, or the direction of maximum change in gene expression within each spatial domain. The collection of spatial gradients defines a spatial transcriptomic vector field v (*x, y*) across the tissue slice *T* (Figure 1A). Thus, the isodepth describes the *geometry* of a tissue slice, i.e. the arrangement of distinct spatial domains in the tissue, as well as directions of continuous variation within each spatial domain (Methods).

**Figure 1:**
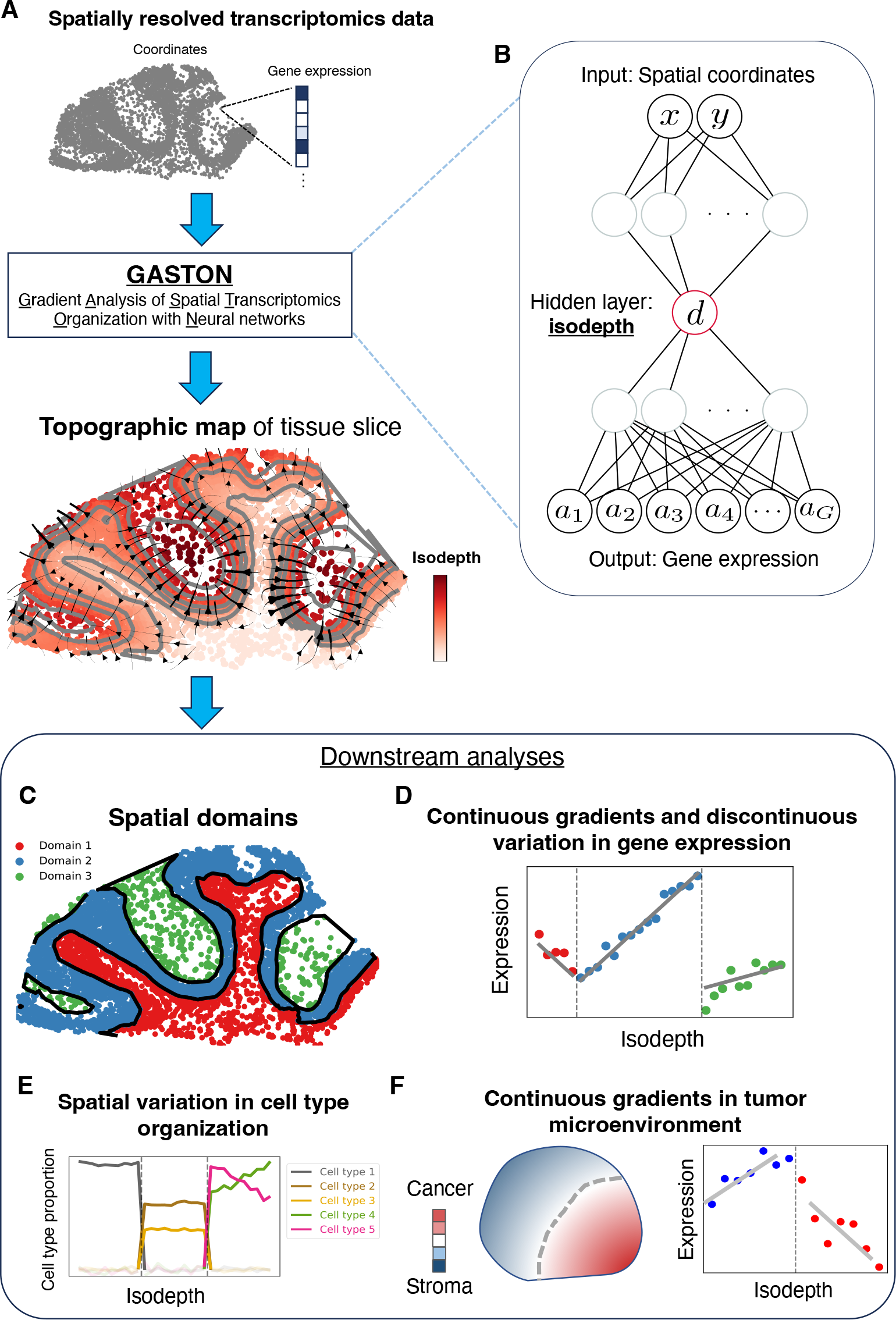
GASTON, an interpretable deep neural network, learns the topography of a tissue. **(A)** GASTON takes in spatially resolved transcriptomics (SRT) data from a tissue slice and outputs the *isodepth*, a coordinate describing a *topographic map* of the tissue slice, with contours of constant isodepth in gray and spatial gradients shown as streamlines. **(B)** GASTON trains a deep neural network to predict gene expression from spatial coordinates, where the isodepth is the value of an *interpretable* hidden layer of the trained neural network. The isodepth learned by GASTON enables many downstream tasks including: **(C)** identification of *spatial domains*, or tissue regions characterized by different cell type composition and gene expression patterns; **(D)** identification of genes with continuous gradients and/or discontinuous variation in expression as a function of isodepth; **(E)** modeling of variation in cell type composition as a function of isodepth; and **(F)** analysis of continuous gene expression gradients in the tumor microenvironment.

To learn the isodepth *d* from spatially resolved transcriptomics (SRT) data, we develop Gradient Analysis of Spatial Transcriptomics Organization with Neural networks (GASTON). GASTON models the expression *f*_*g*_ (*x, y*) of each gene *g* at spatial location (*x, y*) as a *piecewise linear* function of the isodepth *d* (*x, y*):

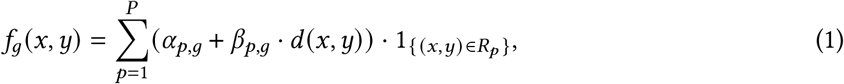

where the pieces *R*_1_, ·, *R*_*P*_ are spatial domains, and α_*p,g*_ and β_*p,g*_ are the *y*-intercept and slope, respectively, in the *p*th spatial domain *R*_*p*_. We use piecewise linear functions as they are a simple class of models that incorporates both continuous variation in gene expression within each domain, i.e. “gradients” of expression, while allowing for discontinuities in expression at the boundaries of the spatial domains. The boundaries of each spatial domain *R*_*p*_ are given by contours of equal isodepth *d* (*x, y*) (Methods). We emphasize that our model does not restrict the spatial domains *R*_*p*_ to be contiguous regions; thus, GASTON is able to model *long-range* spatial correlations in gene expression [101], in contrast to many existing approaches that only model local spatial correlations (Methods).

GASTON jointly learns the isodepth *d* and piecewise linear gene expression functions *f*_*g*_ in a completely unsupervised manner using an interpretable deep learning model. Specifically, GASTON trains a neural network to learn a composite function *f*º*d* (*x, y*) from spatial coordinates to gene expression features, where the isodepth *d* (*x, y*) corresponds to an interpretable hidden layer of the network (Figure 1B). GASTON then uses segmented regression [83, 3, 7] to learn the spatial domains *R*_*p*_, as well as the parameters α, β of the piecewise linear expression functions *f*_*g*_ for each gene *g*. We demonstrate below that GASTON’s interpretable approach uncovers meaningful spatial domains (Figure 1C), and continuous gradients and discontinuities in gene expression (Figure 1D) and cell type composition (Figure 1E) across a wide range of SRT technologies and biological systems including the brain and the tumor microenvironment (Figure 1F).

### 2.2 GASTON recapitulates spatial organization in mouse and human brain slices

We first used GASTON to learn the isodepth *d* and the spatial gradients ∇*d* in a tissue slice from the mouse cerebellum where the expression of 23,096 transcripts at 9,985 spatial locations was measured using the Slide-SeqV2 platform [116, 127]. The learned isodepth *d* provides a “topographical map” of the layered geometry of the cerebellum, including the boundaries of distinct layers of the cerebellum, with the depth within each layer scaled to approximate μm (Figure 2A, Methods). The spatial expression gradients ∇*d* are perpendicular to the cerebellar layers (contours of constant isodepth) and indicate the spatial direction of maximum change in gene expression.

**Figure 2:**
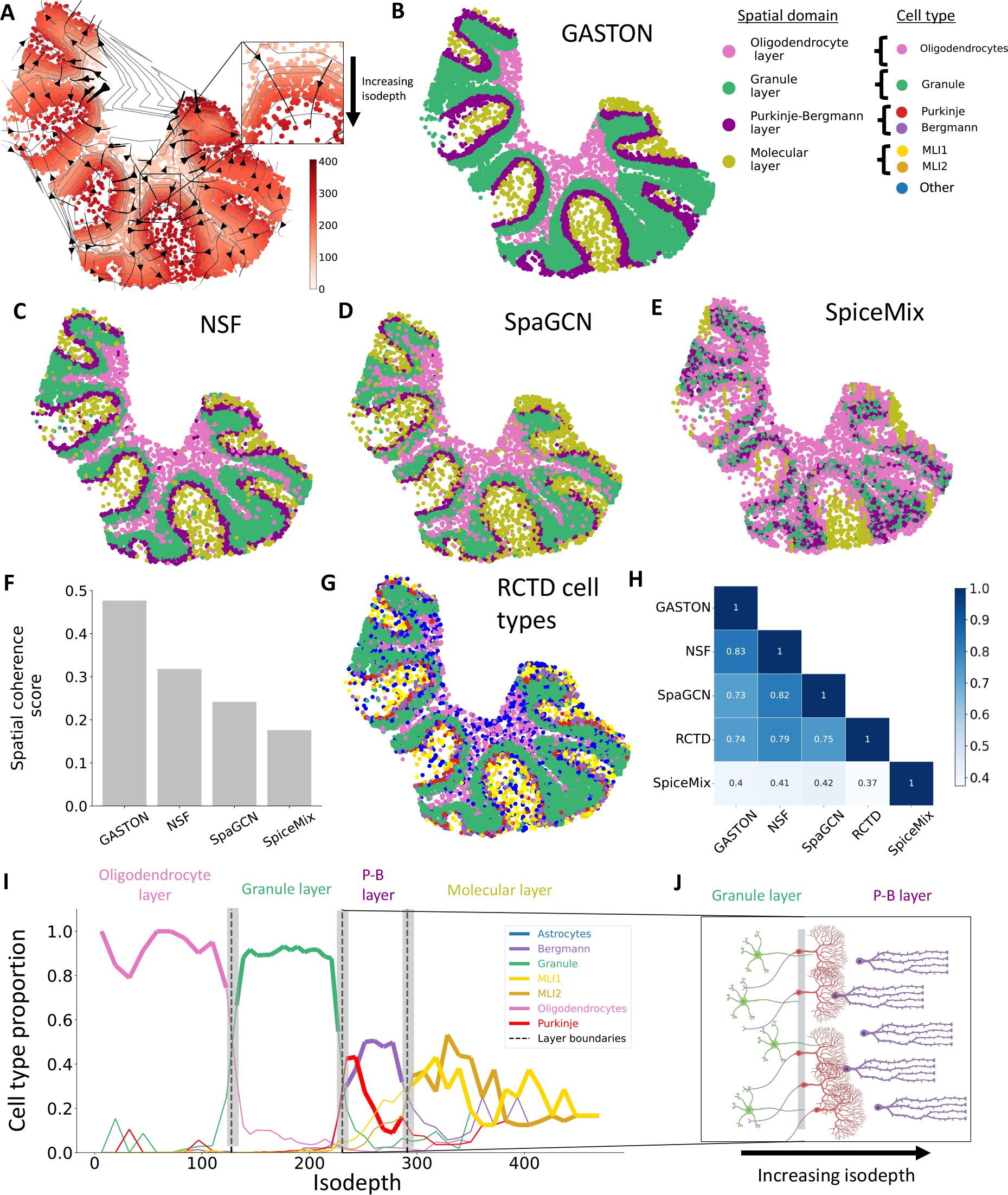
Spatial gradients learned by GASTON recapitulate the spatial organization of the mouse cerebellum. **(A)** The isodepth *d x, y* and spatial expression gradients *d*, shown as streamlines, learned by GASTON on Slide-SeqV2 data from the mouse cerebellum [18]. Gray curves denote contours of equal isodepth. **(B-E)** Spatial domains (layers) *R*_1_, …, *R*_4_ identified using **(B)** GASTON, **(C)** Non-negative Spatial Factorization (NSF), **(D)** SpaGCN, and **(E)** SpiceMix. The spatial domains are colored according to the most prevalent RCTD cell types in the domain. **(F)** Spatial coherence score of spatial domains identified by each method. **(G)** Layer-specific cell types identified by RCTD. **(H)** F-measure between spatial domains identified by GASTON, NSF, SpaGCN, SpiceMix, and layer-specific cell types identified by RCTD. **(I)** Proportions of layer-specific cell types as a function of the isodepth *d*. Dashed lines indicate boundaries of GASTON spatial domains. **(J)** Layout of granule (green), Purkinje (red), and Bergmann (purple) cells as a function of isodepth near the Purkinje-Bergmann layer of the cerebellum.

GASTON divides the tissue into four contiguous spatial domains, which are visually consistent with the four distinct layers of the cerebellum – the oligodendrocyte layer, the granular layer, the Purkinje-Bergmann layer, and the molecular layer – that were identified in prior imaging studies [116, 26] and SRT analyses [116, 19, 18] (Figure 2B). We compared the spatial domains learned by GASTON to those identified by Non-negative Spatial Factorization (NSF) [135], SpaGCN [58], and SpiceMix [24] (Figure 2C-E), three recent methods that showcase the major approaches currently used to model local spatial correlations in spatial transcriptomics data: Gaussian processes (GPs), graph convolutional networks (GCNs), and hidden Markov random fields (HMRFs), respectively. We observed that GASTON’s spatial domains have much larger spatial coherence [159] compared to the other methods (Figure 2F), showing that the domains identified by GASTON better align with the structured geometry of the cerebellum [140]. Next, we compared the spatial domains to the cell types reported in the original publication of the data (Figure 2G). These cell types were obtained from RCTD [19], a method which performs cell type deconvolution using a reference scRNA-seq dataset and does not take spatial information into account. The GASTON, SpaGCN, and NSF spatial domains have similar agreement with the cell types inferred by RCTD and with each other, while the SpiceMix spatial domains have low agreement with the RCTD cell types and the other methods (Figure 2H). These results demonstrate that the global model of spatial variation used in GASTON identifies more spatially coherent spatial domains than existing methods while still preserving cell type information.

A key distinguishing feature of GASTON is that it learns the isodepth *d*, which provides a coordinate to analyze the continuous variation in cell types within and across the layers of the cerebellum. Such continuous variation is not modeled by the three methods above nor by the numerous other methods that divide a tissue slice into spatial domains, e.g. [168, 32]. We find that the proportion of cell types varies considerably as a function of the isodepth (Figure 2I). First, we observe that oligodendrocytes and granule cells have large and nearly constant proportion throughout the range of isodepth *d* that corresponds to the named layers. Moreover, there is a sharp transition in proportion at the isodepth value that GASTON marks as the boundary between these layers, indicating that the learned isodepth *d* and spatial domains are accurately separating the oligodendrocyte and granule layers.

In contrast, the proportion of Purkinje cells and Bergmann glia exhibit spatial variation with the Purkinje-Bergmann layer. Purkinje cells are concentrated at the start of the layer (small isodepth), while the Bergmann glia peak in proportion inside the layer and are present over a wider range of isodepths (Figure 2J). These results agree with prior imaging and microscopy-based studies which show that Purkinje cells form a “monolayer” in the cerebellum, i.e. a layer with single-cell depth [170, 126, 13] while the Bergmann glia do not form a monolayer but are more diffusely spread out across the Purkinje-Bergmann layer [5, 72]. Interestingly, previous studies have found that the Bergmann glia form a monolayer during the development of the cerebellum [68, 54], and thus the observed arrangement of Bergmann glia here could indicate that the spatial arrangement of Bergmann glia changes after development. We also observe that the Bergmann glia are closer to the molecular layer of the cerebellum compared to Purkinje cells, which agrees with previous studies on cerebellar organization [115].

We emphasize that GASTON learns the isodepth *de novo* and in an unsupervised manner. In contrast, existing approaches for learning depth or depth-like measurements either require prior anatomical knowledge [83, 84], which is difficult to obtain for a complex tissue like the cerebellum, or use scRNA-seq-based trajectory inference approaches which do not learn a spatially continuous measurement (see comparison to SpaceFlow [113] in Supplement C, Figure S1).

As additional validation, we evaluated GASTON using SRT data of the human dorsolateral prefrontal cortex (DLPFC) [89]. GASTON more accurately identified the manually annotated layers of the DLPFC compared to two graph neural network approaches: SpaGCN [58] and STAGATE [32] (Figure S3). Moreover, GASTON has comparable performance to Belayer [83], which previously achieved state-of-the-art performance in DLPFC layer identification using prior annotation on the layer boundaries. In contrast, GASTON, an unsupervised algorithm, achieves similar performance without any no prior knowledge. See Supplement D for details.

These analyses demonstrate that the isodepth *d* learned by GASTON provides a powerful computational approach for modeling the spatial organization of cells and cell types in complex biological tissues.

### 2.3 Continuous and discontinuous spatial variation in gene expression

We next investigated whether GASTON identifies biologically meaningful spatial patterns of gene expression in sparse SRT data, particularly in low coverage Slide-SeqV2 data (median≈500 UMIs per spatial location [127]) where such patterns may not be apparent. For each gene *g*, GASTON learns a piecewise linear function *h*_*g*_(*d*) of the isodepth *d* that models both continuous variation in expression within or across spatial domains and sharp discontinuities in gene expression between adjacent spatial domains. These learned gene expression functions (Supplementary Table) indicate genes that have spatially varying expression patterns. For example, *SBK1* – reported to be a marker gene of Purkinje cells [71] – has particularly sparse expression in the Slide-SeqV2 cerebellum tissue, with only 15% of all spatial locations having non-zero UMI count, and only 2% of spatial locations in the GASTON-estimated Purkinje-Bergmann layer having UMI count *>* 1. (Figure 3A). By aggregating expression across contours of constant isodepth (Figure 2A), GASTON learns a piecewise linear gene expression function for *SBK1* that peaks in the Purkinje-Bergmann layer and exhibits continuous variation in the granule layer as a function of isodepth (Figure 3B). The corresponding 2D expression function clearly demarcates the Purkinje-Bergmann layer (Figure 3C) compared to the sparse expression values (Figure 3A).

**Figure 3:**
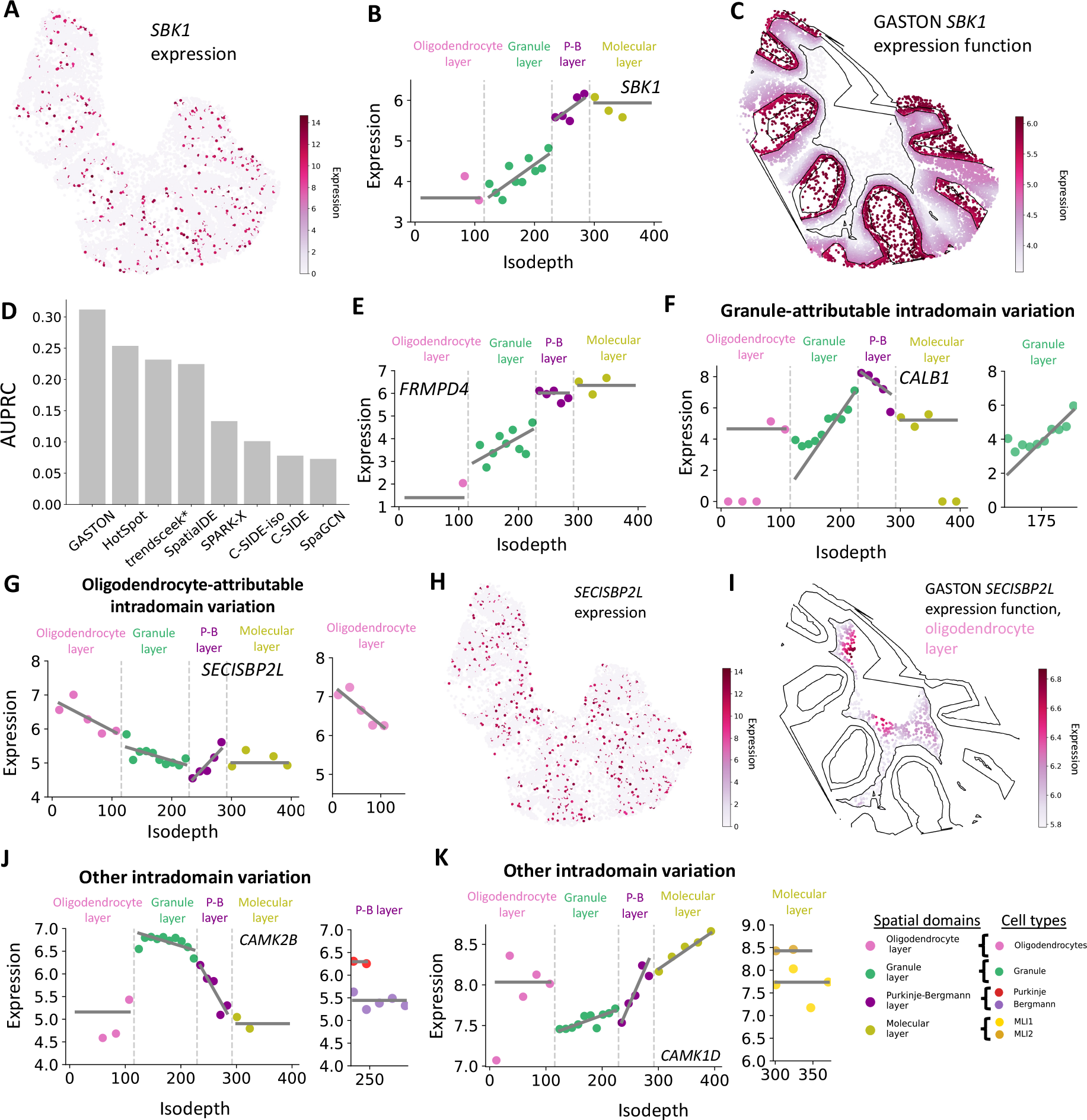
GASTON reveals continuous and discontinuous spatial variation in gene expression in the mouse cerebellum. **(A)** *SBK1* expression, shown in log counts per million (CPM). **(B)** Isodepth versus expression for *SBK1*. Lines denote piecewise linear function *h*_*g*_ *d* learned by GASTON. **(C)** *SBK1* expression function *f x, y* learned by GASTON. Curves denote contours of constant isodepth *d*. **(D)** Comparison of GASTON and several existing methods in marker gene identification, quantified using the area under precision-recall curve (AUPRC) and a list of known cerebellum marker genes [40, 71, 118, 69]. trend-sceek* uses the Seurat [50] implementation and C-SIDE-iso runs C-SIDE using the isodepth *d* learned by GASTON as a covariate. **(E)** Isodepth versus expression for *FRMPD4* which was ranked highly by GASTON as a marker gene in (D). **(F)** (Left) Isodepth versus expression for *CALB1*, which has (Right) granule-attributable intradomain variation since the expression function restricted to granule cells has large slope. **(G)** (Left) Isodepth versus expression for *SECISBP2L* which has (Right) oligodendrocyte-attributable intradomain variation since the expression function restricted to oligodendrocyte cells has large slope. **(H)** *SECISBP2L* expression shown in log CPM. **(I)** *SECISBP2L* expression function *f x, y* learned by GASTON in the GASTON-inferred oligodendrocyte layer. **(J-K)** Isodepth versus expression for **(J)** *CAMK2B* and **(K)** *CAMK1D* which have (Left) intradomain variation in the Purkinje-Bergmann layer and molecular layer, respectively, that is (Right) not attributable to cell type, as the expression functions for the most abundant cell types in the respective layers have zero slope.

The gene expression functions learned by GASTON yield a substantially better predictor of known marker genes in the cerebellum than existing methods for identifying spatially variable genes (SVGs) or differentially expressed genes (DEGs). Specifically, by ranking genes according to a measure of the variance of the GASTON expression function across spatial domains (Methods), GASTON achieved notably higher performance (AUPRC≈0.31) in the identification of marker genes compared to HotSpot [31]; trendsceek [35]; SpatialDE [132]; SPARK-X [130, 171]; C-SIDE [18]; and SpaGCN [58] which have AUPRC ranging from 0.07 to 0.25 (Figure 3D). A major reason for GASTON’s improved performance is because many of the other methods test only whether the expression of each gene varies in 2D space, and are unable to distinguish between different types of continuous and discontinuous variation in spatial expression. In contrast, GASTON’s piecewise linear gene expression function explicitly models both continuous and discontinuous variation in expression. We highlight two genes ranked highly by GASTON but not by other methods: *SBK1*, described previously, and *FRMPD4. FRMPD4* is not a known marker gene but has high expression in the molecular layer (Figure 3E). Recent studies report that the *FRMPD4* protein regulates neurons in the molecular layer, with mutations of *FRMPD4* causing intellectual disabilities [105].

As another demonstration of the utility of the isodepth *d* learned by GASTON, we used the isodepth as a covariate for C-SIDE [18], which identifies cell type-specific differentially expressed (DE) genes from SRT data. This variation of C-SIDE, which we call C-SIDE-iso, identifies a substantially different set of DE genes compared to the original C-SIDE, with only a 10% overlap between the DE genes identified by both approaches. C-SIDE-iso achieved better performance than the original C-SIDE in marker gene identification (Figure 3C), demonstrating the advantages of the isodepth *d*. Nevertheless, unlike GASTON, C-SIDE-iso cannot identify spatial domains and thus cannot test for changes in expression across different spatial domains, and consequently C-SIDE-iso has lower performance than GASTON in identification of marker genes (Figure 3C).

In addition to marker gene identification, the piecewise linear expression functions learned by GASTON reveal distinct spatial patterns of gene expression including discontinuities in expression — i.e. large differences in expression between adjacent spatial domains — or continuous *intradomain* variation — i.e. a large slope β of the piecewise linear expression function within a spatial domain (Methods). GASTON identifies 513 spatially varying genes with either discontinuities or intradomain variation (Figure S2A). Approximately half of these genes have discontinuities in expression, indicating that a gene is selectively expressed or not expressed within cells in a specific spatial domain. For example, GASTON finds that *CPLX2* has discontinuities in expression at the boundaries of the granule layer, which matches prior studies showing that large expression of *CPLX2* in granule cells suppresses differentiation pathways [155] (Figure S2B). Furthermore, more than 60% of the spatially varying genes identified by GASTON have continuous intradomain variation (Figure S2A), indicating that continuous variation is fairly common in the cerebellum. This observation may explain why SpaGCN, whose clustering algorithm assumes there is no continuous variation in gene expression, is less accurate in resolving the layers of cerebellum layers (Figure 2D).

Continuous intradomain variation in gene expression may be due to a continuum of cell states within a cell type, continuous variation in the proportion of cell types in a tissue, or other causes [160]. We evaluated whether the intradomain variation identified by GASTON was attributable to the annotated cell types in each domain, which distinguishes whether there is (1) a spatial component in the continuum of cell states within a cell type [160] or (2) spatial variation in either the proportion of cell types or other causes (Methods). Specifically, we say that intradomain variation is *cell type-attributable* if the slope β_*c*_ estimated only from cells annotated as cell type *c* has magnitude |β_*c*_| close to or larger than the magnitude |β| of the slope β estimated from all cells (Methods). We find that 217 of the 338 genes that GASTON reports to have intradomain variation have cell type-attributable intradomain variation (Figure S2A).

The cell type-attributable intradomain variation identified by GASTON reveals important cell type-specific processes including neuronal firing and differentiation. For example, *CALB1*, which is involved in calcium binding, has granule-attributable intradomain variation in the granule layer (Figure 3F). This granule-attributable *CALB1* continuous variation identified by GASTON provides a potential molecular explanation for the reported spatial gradients in neuronal firing thresholds for granule cells in the granular layer [128]. A second example is *SECISBP2L*, which exhibits large oligodendrocyte-attributable intradomain variation in the oligodendrocyte layer (Figure 3G). *SECISBP2L* was recently shown to be specifically expressed in differentiating oligodendrocytes, with *SECISBP2L* more highly expressed in less mature oligodendrocyte cells [29]. The observed decrease in *SECISBP2L* expression as a function of isodepth suggests that oligodendrocyte differentiation may occur along the isodepth axis, i.e. along the spatial gradients ∇*d*, in the oligodendrocyte layer (Figure 2A). Notably, continuous variation in *SECISBP2L* expression in the oligodendrocyte layer is not apparent from individual expression values per spot (Figure 3H), but is revealed by the expression function learned by GASTON, which pools expression values along contours of constant isodepth (Figure 3I).

Approximately 35% of the intradomain variation in gene expression identified by GASTON is not attributable to cell type (Figure S2A). For example, *CAMK2B*, which is overexpressed in granule cells, has large intradomain variation in the Purkinje-Bergmann layer (Figure 3J, left). However, this intradomain variation is not attributable to the Purkinje or Bergmann cell types, as the Purkinje- and Bergmann-specific expression functions for *CAMK2B* have zero slope (Figure 3J, right). Instead, the intradomain variation of *CAMK2B* is likely attributable to the large decrease in *proportion* of granule cells in the Purkinje-Bergmann layer as a function of isodepth (Figure 2I). *CAMK1D*, a calcium-dependent protein kinase whose aberrant behavior has been linked to Alzheimer’s disease [47] and glioma [63], exhibits intradomain variation in the molecular layer (Figure 3I, left) that is not attributable to either MLI1 or MLI2 neurons (Figure 3I, right). This variation could be attributable to other causes such as cellular interactions or neuronal firing.

These analyses demonstrate that GASTON’s combined model of continuous and discontinuous variation of gene expression reveals biologically meaningful marker genes and continuous gradients of expression not found by existing approaches.

### 2.4 Spatial gradients in the tumor microenvironment

We next used GASTON to investigate spatial gene expression patterns in the tumor microenvironment (TME). The TME is strongly correlated with tumor development and prognosis [43], but is challenging to quantify accurately without spatial information [157]. However, existing analyses of tumor SRT data, e.g. [12, 36, 62], examine only differentially expressed (DE) genes or pathways between the tumor and surrounding stromal regions. We hypothesized that GASTON’s ability to quantify continuous variation might reveal more subtle variation in gene expression relative to the boundary of the tumor.

We applied GASTON to SRT data from a colorectal (CRC) tumor tissue slice (Figure 4A) where the expression of 36,601 transcripts in 3,900 spots was measured using the 10x Genomics Visium platform [149]. GASTON identifies five spatial domains (Figure 4B) that are visually distinct in the H&E-stained image (Figure 4A), including the tumor (domain 1), the tumor-adjacent stromal region (domain 2), and other stromal regions not directly adjacent to the tumor (domains 3-5). In contrast, the the published analysis of this data performed unsupervised clustering of spots based on gene expression alone [121] and was unable to distinguish between the different stromal regions of the the tissue slice (Figure S4).

**Figure 4:**
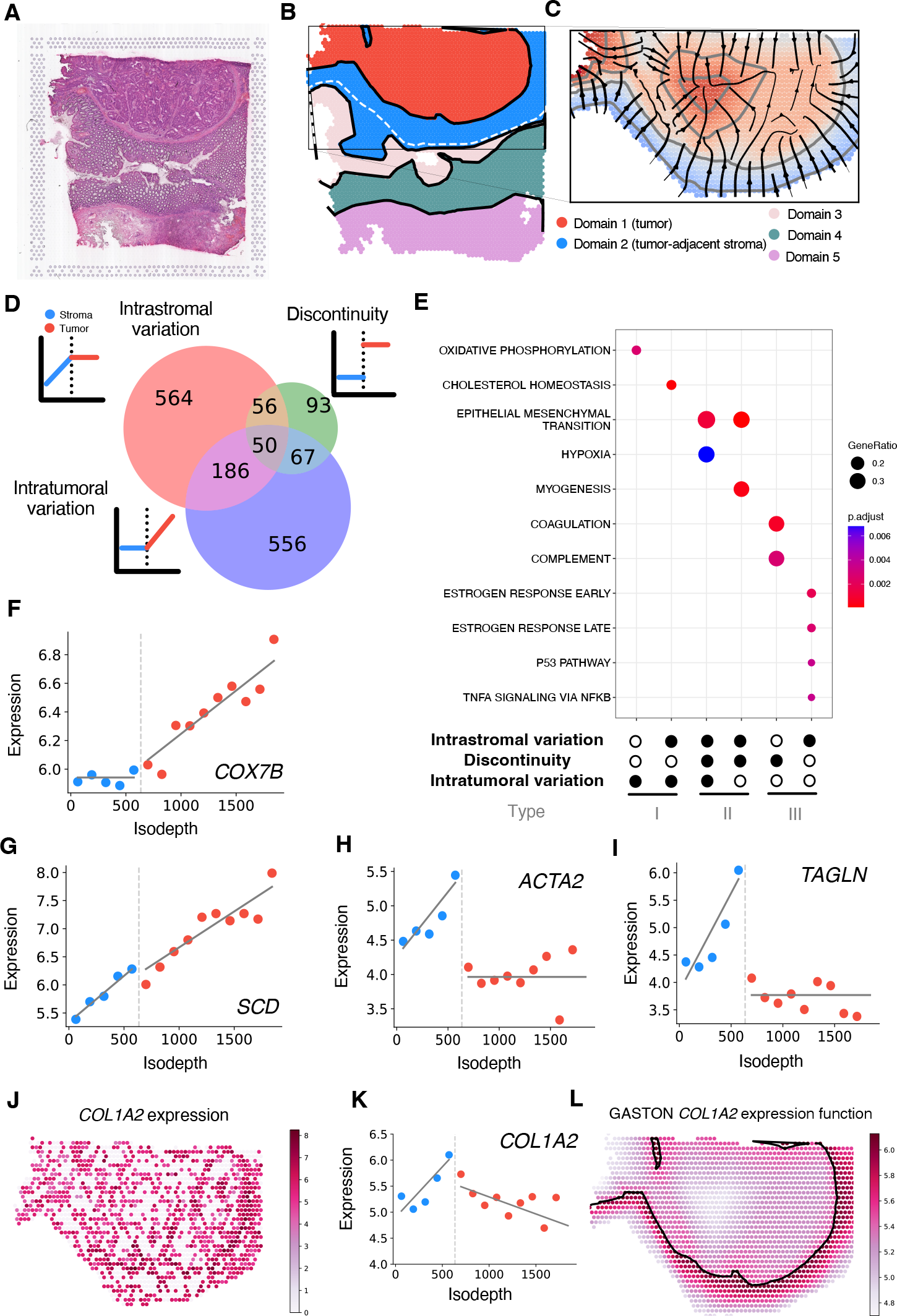
GASTON identifies spatial gene expression patterns in the tumor micro-environment. **(A)** H&E stain of a 10x Genomics Visium colorectal tumor sample. **(B)** Spatial domains learned by GASTON. Domains 1 and 2 are labeled as tumor and tumor-adjacent stroma, respectively, based on the histology image in (A). **(C)** Isodepth *d* and spatial gradients learned by GASTON restricted to tumor and tumor-adjacent stromal domains. **(D)** GASTON identifies 986 spatially varying genes which are classified into three spatial expression patterns: genes with intrastromal variation in expression; genes with a discontinuity in expression at the tumor-stroma boundary; and genes with intratumoral variation in expression. **(E)** Enrichment for hallmark cancer gene sets reported by gene set enrichment analysis (GSEA) for six of the seven spatial expression patterns in (D). The spatial expression patterns are grouped into three types according to expression pattern and enriched cancer pathways. **(F-I)** Isodepth *d* versus expression for Type I genes **(F)** *COX4l1* and **(G)** *SCD*, and Type II genes **(H)** *ACTA2* and **(I)** *TAGLN*. **(J)** *COL1A2* expression shown in log CPM. **(K)** Expression versus isodepth for Type II gene *COL1A2*. **(L)** GASTON *COL1A2* expression function shows a gradient of expression at the tumor-stroma boundary.

We analyzed spatial variation in the TME by examining the expression of each gene as a function of the isodepth *d*, which varies smoothly from the tumor boundary to the interior (Figure 4C, Supplementary Table). GASTON identifies 1,572 spatially varying genes in the tumor and adjacent stromal domains which exhibit one of seven different spatial expression patterns: intratumoral variation, a discontinuity at the tumor-stroma boundary, intrastromal variation, or any combination of these (Figure 4D). For six of the seven spatial gene expression patterns, the genes exhibiting the spatial pattern are enriched (*p <* 0.01, GSEA [74]) for cancer hallmark gene sets (Figure 4E). We further group the genes in the six enriched spatial gene expression patterns found by GASTON into three different types: (1) Type I genes, which have intratumoral variation and no discontinuity in expression; (2) Type II genes, which have intrastromal variation and a discontinuity at the tumor-stroma boundary; and (3) Type III genes, which have either intrastromal variation or discontinuity at the tumor-stroma boundary but no intratumoral variation.

The three types of spatially varying genes identified by GASTON reflect distinct biological processes occurring in the TME. The 742 Type I genes (intratumoral variation) are enriched for oxidative phosphorylation and cholesterol homeostasis gene sets; moreover, 39 of the 42 Type I genes involved in oxidative phosphorylation or cholesterol homeostasis have *positive* slopes within the tumor domain, indicating an increase in expression from the margin to the interior of the tumor. Thus, Type I genes likely indicate an *increasing* gradient of metabolic activity from the tumor boundary to the interior [74]. For example, *COX7B* (Figure 4F) is a Type I gene in the oxidative phosphorylation pathway and a component of the cytochrome c oxidase protein complex which transfers electrons to oxygen in the electron transport chain and leads to ATP synthesis [144]. Several other genes in this complex are also Type I genes, including *COX17, COX7A2, COX6C*, and *COX8A*. Another Type I gene is Stearyl-CoA desaturase (*SCD*, Figure 4G), a fatty enzyme that is key component of lipid metabolism [122], with *SCD* deficiency being linked to reduced lipid synthesis and other poor health outcomes [38]. Interestingly, the expression of both *SCD* and *COX7B* are directly affected by oxygen availability [151], with lower expression in hypoxic conditions. The higher expression of these genes in the interior of the tumor suggests that the interior of this CRC tumor slice is more oxygenated than the boundary. This observation is consistent with a previous clinical study which found that that stage IV CRC tumors may have lower hypoxia response — and thus higher oxygen availability — in the tumor interior compared to the boundary [2].

The 106 Type II genes (intrastromal variation and discontinuity) primarily describe the upregulation of epithelial-mesenchymal transition (EMT) genes immediately outside the tumor boundary. Several studies have shown that upregulation of EMT genes within tumor-associated stromal cells is associated with aggressive, poor prognosis CRC subtypes [20, 60, 73]. Of the 15 type II genes in the EMT pathway, 14 had positive slopes with isodepth in the tumor-adjacent stroma domain, i.e. expression increased closer to the tumor boundary, suggesting that this stage IV colorectal tumor was likely an aggressive subtype. For example, *ACTA2* and *TAGLN*, two genes that were reported to be markers of a subtype of colorectal cancer-associated fibroblasts with upregulated EMT-related genes [73], have positive slopes and large discontinuities at the tumor boundary (Figure 4H, I). GASTON also finds that *ACTA2* and *TAGLN* have constant, low expression in the tumor region, consistent with previous studies that find no evidence for upregulation of EMT-related genes in CRC tumor cells [20, 60]. The upregulation of EMT genes – such as *ACTA2* and *TAGLN* – in tumor-associated stromal cells could be an important mechanism underlying the aggressiveness of this CRC tumor, where these stromal cells may facilitate local invasion and metastasis [65]. Notably, the overexpression of several Type II genes is concentrated on the right side of the tumor boundary (Figure S5A,B), suggesting that the local invasion and metastasis may be localized to a specific part of the tumor boundary. We also highlight the Type II gene *LGR5*, which has large expression at the tumor boundary and has been reported to be a potential marker for CRC stem cells [92] (Figure S5C,D). The co-expression of *LGR5* and *ACTA2* / *TAGLN* suggests a potential interaction between tumor-adjacent stromal cells and CRC stem cells.

We emphasize that the upregulation of EMT genes near the tumor boundary is not readily apparent from the sparse UMI counts. For example, *COL1A2* is a Type II gene involved in EMT [146], but the spatial distribution of *COL1A2* expression is difficult to discern directly (Figure 4J), with nearly half of all spots having no measured *COL1A2* transcripts while only a small fraction of spots (5%) have more than 10 transcripts. GASTON aggregates the sparse *COL1A2* expression measurements across the contours of constant isodepth and learns a piecewise linear *COL1A2* expression function of isodepth (Figure 4K), revealing continuous variation in *COL1A2* expression. In particular, GASTON finds that *COL1A2* expression peaks at the tumor boundary and decays in the interior of tumor and in the tumor-adjacent stroma (Figure 4L). This expression pattern is consistent with a recent report demonstrating that *COL1A2* expression is lower in primary CRC tumors compared to adjacent stromal tissue [156].

The 657 Type III genes (no intratumoral variation) identified by GASTON primarily describe immune response in the stroma as well as cell signaling and proliferation in the tumor. For example, *THBS1* has a large discontinuity in expression at the tumor-stroma boundary and has high expression in the tumor-adjacent stroma (Figure S5E), consistent with reports that *THBS1* expression promotes immune cell response in other cancer types [106, 166]. Another Type III gene, *FUCA1*, is involved in fucosylation of proteins and a member of the p53 signaling pathway [37]. GASTON finds that *FUCA1* has large, negative slope in the stroma region; no discontinuity in expression at the tumor boundary; and constant, low expression in the tumor region (Figure S5F). This spatial expression pattern suggests that *FUCA1* is downregulated in the tumor region, agreeing with several recent studies which found that *FUCA1* is downregulated in highly aggressive and metastatic CRC and breast tumors [16, 99].

Overall, the spatial gene expression patterns identified by GASTON suggest that the interior of this CRC tumor sample is growing slowly – since aerobic metabolism through oxidative phosphorylation indicates slow cellular growth and proliferation [169] – while the boundary is undergoing EMT to stem-like states [86]. These features of the tumor interior and boundary indicate a late-stage, vascularized primary tumor with a fully metastatic margin, a characterization which aligns with the tumor’s clinical information [149]. Thus, this analysis demonstrates how the gene expression topography learned by GASTON enables the characterization of the spatial and molecular organization of the TME.

### 2.5 Spatial gradients of cell type and gene expression in the mouse olfactory bulb

Finally, we used GASTON to analyze Stereo-seq [22] data from the mouse olfactory bulb which measures the expression of 27,106 transcripts at 9,825 spatial locations. Stereo-seq achieves single cell spatial resolution using DNA nanoball patterned array chips, but the data is highly sparse, with a median UMI of less than 350 per location. At the same time, the olfactory bulb has a *radial* geometry consisting of several concentric layers (Figure 5A), and this geometry provides spatial constraints that may help overcome the severe data sparsity.

**Figure 5:**
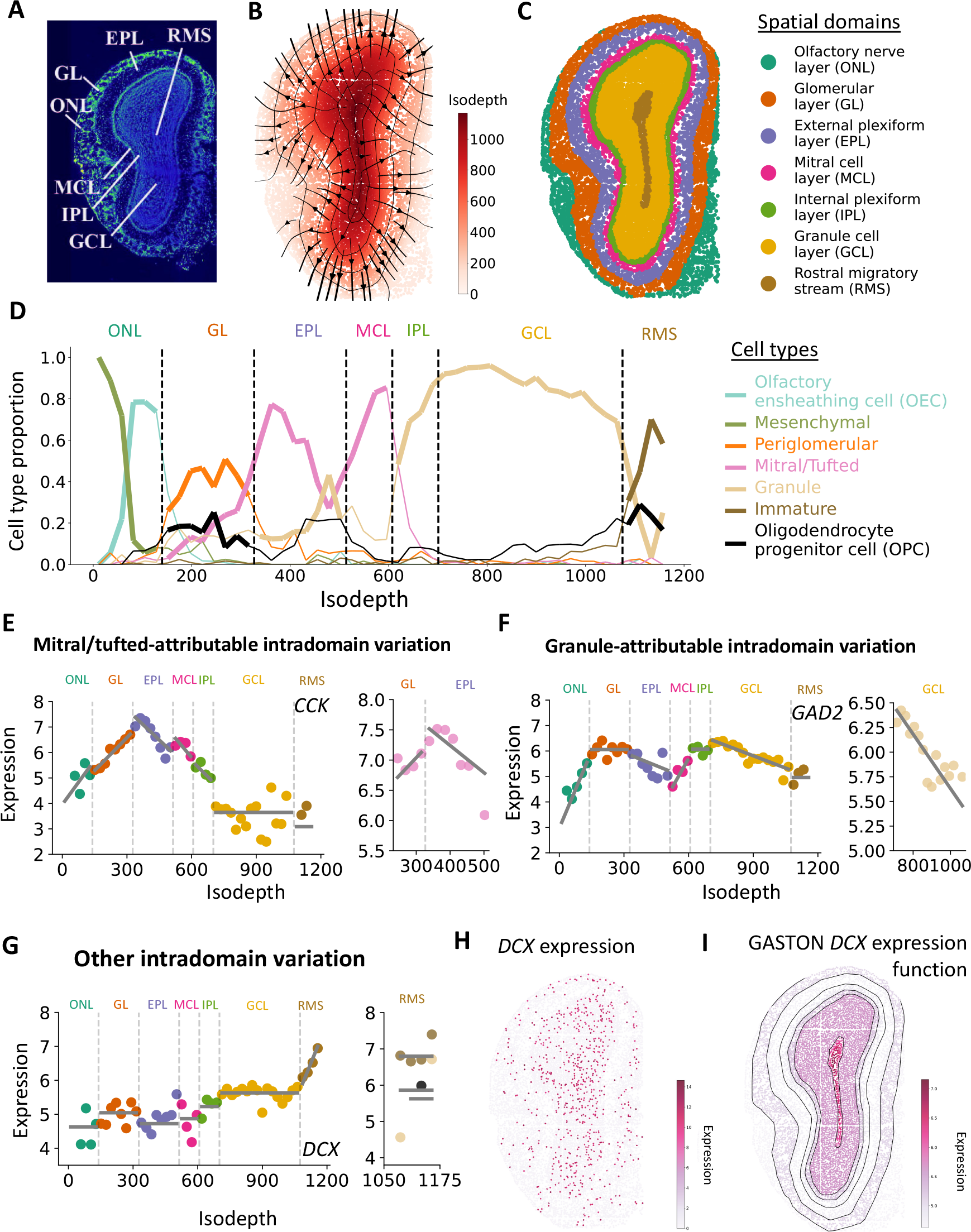
GASTON reveals variation in cell types and gene expression in the mouse olfactory bulb. **(A)** DAPI stain of mouse olfactory bulb [42] produced by [22]. **(B)** Isodepth *d* and (negative) spatial gradients −∇*d* (shown as streamlines) learned by GASTON. Curves denote contours of constant isodepth *d*.**(C)** Spatial domains learned by GASTON and labeled based on annotations in (A). **(D)** Cell type proportion as a function of isodepth *d*. Dashed lines indicate boundaries of spatial domains identified by GASTON. Most abundant cell types in each spatial domain are highlighted. **(E)** (Left) Isodepth versus expression for *CCK* which (Right) has mitral/tufted-attributable intradomain variation in the glomerular layer (GL) and external plexiform layer (EPL). **(F)** (Left) Isodepth versus expression for *GAD2* which (Right) has granule-attributable intradomain variation in the granule cell layer (GCL). **(G)** (Left) Isodepth versus expression for *DCX* which has continuous variation in the rostral migratory stream (RMS) which is (Right) not attributable to cell type, as the expression function for the most abundant cell types have zero slope. **(H)** *DCX* expression shown in log CPM. **(I)** *DCX* expression function learned by GASTON.

GASTON learns the radial geometry of the olfactory bulb nearly perfectly, with the isodepth *d* providing a topographic map that reflects the geometry of the olfactory bulb (Figure 5B). Using the learned isodepth, GASTON divides the tissue into seven contiguous spatial domains (Figure 5C) that visually correspond to the seven distinct layers of the olfactory bulb (Figure 5A). In comparison, the spatial domains found by SpaGCN, a method based on a graph convolutional neural network, are less spatially coherent than the GASTON domains and do not reflect the layered geometry of the olfactory bulb. Notably, SpaGCN poorly resolves the innermost rostral migratory stream (RMS) layer (Figure S6A,B).

The olfactory bulb is one of two regions in the brain where adult neurogenesis occurs, with immature neurons migrating outward from the RMS (large isodepth) towards the outermost olfactory nerve layer (ONL, small isodepth) [90, 80]. Thus, in this tissue, the isodepth *d* learned by GASTON provides a measure of *potency* in the olfactory bulb, and the negative gradients −∇*d* show the spatial trajectories of neural maturation and migration (Figure 5B).

GASTON reveals substantial variation in cell type composition as a function of isodepth *d* in the olfactory bulb (Figure 5D). While the cell type composition of the different layers of the olfactory bulb is well-studied, GASTON uncovers the spatial arrangement of cell types within each layer which has not been fully characterized in the literature [94]. For example, while previous studies have found that both mesenchymal cells and olfactory ensheathing cells (OECs) are in the outermost olfactory nerve layer (ONL) [75], GASTON identifies that these two cell types have different spatial arrangements in the ONL: mesenchymal cells are concentrated on the outer edge of the layer (isodepth*d <* 50) while OECs peak at a larger isodepth (*d*≈85) and are spread more diffusely throughout the ONL. This arrangement of mesenchymal cells aligns with studies showing that ONL neuron axons grow towards mesenchymal cells during development [33, 109], as axons in the olfactory bulb point outwards [75], i.e. towards small isodepth. In the interior of the olfactory bulb, GASTON finds that immature neurons are most prevalent in the rostral migratory stream (RMS), with the proportion of immature neurons increasing sharply with isodepth, in agreement with studies showing that neurogenesis occurs starting from the RMS interior [90, 80].

The isodepth *d* also distinguishes different cell types or cell states with similar gene expression profiles. For example, mitral cells and tufted cells are grouped together in the single-cell reference dataset [133] used for cell type annotation, and also by SpaGCN (Figure S6A), due to the similar gene expression profiles of these cell types. However, GASTON reveals that the proportion of mitral/tufted cells peaks at two different isodepth values, *d*≈350 and *d*≈600, with a larger proportion of mitral/tufted cells at the second peak versus the first peak (Figure 5D). This suggests that the mitral/tufted cells at isodepth *d*≈350 are tufted cells, which previous studies have shown are spread diffusely in the external plexiform layer (EPL) [94], while the mitral/tufted cells at isodepth *d*≈600 are mitral cells, which have been shown to form a monolayer in the mitral cell layer (MCL) [41]. GASTON also distinguishes between different granule cell states. While there is a single category of granule cells in the single-cell reference dataset [133], previous studies have shown that there are morphologically distinct granule cell subtypes in different layers of the olfactory bulb [94]. GASTON shows that while granule cells are most prevalent in the granule cell layer (GCL), there is a small population of granule cells in the EPL and in the MCL with roughly constant cell type proportion (Figure 5D). The spatial segregration of these two granule cell populations suggests that the granule cells in the EPL and MCL may have a different cell state compared to granule cells in the GCL. Notably, neither the distinction between mitral and tufted cells nor the prevalence of immature neurons in the interior of the bulb are apparent using an alternative 1-D coordinate computed by SpaceFlow [113] that is based on diffusion pseudotime [49] (Supplementary Section C and Figure S6).

GASTON identifies 704 genes with spatially varying expression – i.e. genes with either discontinuous expression or intradomain variation in expression – in the olfactory bulb (Figure S7, Supplementary Table). These genes distinguish different cell types and states in the olfactory bulb and reveal potential molecular mechanisms for biological phenomena. We highlight three examples here. *CCK*, which is reported to be a marker for a specific subtype of tufted cells [163, 131, 59], has mitral/tufted-attributable intradomain variation in the glomerular layer (GL) and EPL (Figure 5E). As noted above, the mitral/tufted cells in the GL and EPL are likely tufted cells, indicating that the continuous variation in *CCK* expression is likely tufted cell-attributable and not mitral cell-attributable. *GAD2*, a marker gene for neurons in the GABAergic systems– the main inhibitory neurotransmitter system in the brain [8, 14, 10] – has granule-attributable intradomain variation in the GCL (Figure 5F). Granule cells are known to be GABAergic [94], suggesting that the granule-attributable variation identified by GASTON may play a role in the GABAergic system. *DCX* (doublecortin) has large intradomain variation in the RMS (Figure 5G), consistent with reports [39, 45] that *DCX* is a marker gene for immature neurons in the RMS (Figure 5D). The continuous variation in *DCX* expression is not attributable to cell type, and instead is likely due to the increasing *proportion* of immature neurons in the RMS as a function of isodepth (Figure 5D). While the intradomain variation in *DCX* expression is challenging to observe from the sparse Stereo-seq UMI counts (Figure 5H), GASTON learns a *DCX* expression function that pools expression across isodepth and uncovers the continuous intradomain variation in *DCX* (Figure 5I).

## 3 Discussion

Accurate models of spatial gene expression variation within tissues are critical for determining the spatial organization of cell types and for defining processes of differentiation and intercellular communication that modulate cell states within spatial niches. Spatial variation in gene expression includes both discontinuous changes in gene expression across the different spatial domains of a tissue, as well as continuous variation within and across spatial domains due to variation in cell state or other causes. While numerous computational methods have been developed to identify spatial domains by modeling discontinuous changes in gene expression, few methods are able to identify spatial domains and *simultaneously* model continuous variation within the domains. Moreover, to our knowledge no existing methods perform this simultaneous identification in an unsupervised and biologically interpretable manner.

In this work, we introduce the *isodepth*, a coordinate that models both the arrangement of spatial domains within a tissue and the relative position of spatial locations with each domain. The isodepth gives a *topographic map* of a tissue slice, analogous to elevation in a map of the Earth’s surface. The gradient of the isodepth describes *spatial gradients*, or the spatial directions of maximum change in gene expression in a tissue. We derive an unsupervised and interpretable deep learning algorithm, GASTON, that learns the isodepth, spatial gradients, and piecewise linear gene expression functions of the isodepth. We demonstrate that the isodepth and spatial gradients learned by GASTON improve detection of spatial domains and spatially varying marker genes, and enable the identification of spatial gene expression patterns linked to important biological processes including differentiation and communication in the brain as well as hypoxia in the tumor microenvironment.

A key advantage of the isodepth computed by GASTON is that it provides a *global* model of spatial gene expression. Just as one can climb to the same elevation on two different mountains, so too can the isodepth take on the same value at two spatially separated locations in the same spatial domain, e.g. the Purkinje-Bergmann layer in the mouse cerebellum (Figure 2B). This global model presents a stark departure from nearly all existing SRT methods which model only *local* spatial correlations. Using the isodepth, GASTON is able to model *“long-range”* spatial correlations, i.e. correlations between distant spatial locations, and pool information across spatially distant locations on the same isodepth contour. As we demonstrate, incorporating these long-range dependencies leads to improved inference of spatial domains and marker genes.

On a smaller scale, the isodepth learned by GASTON provides a coordinate for quantifying variation in gene expression in the tumor microenvironment (TME). Just as single-cell transcriptomics of tumor samples led to the identification of numerous clinical and molecular biomarkers [158], we anticipate that spatial variation in gene expression in the TME will also have high clinical relevance. For example, we showed that GASTON extracts gradients in gene expression that correlate with metabolism, the epithelial-mesenchymal transition (EMT) and other hallmarks of the TME which may translate to novel biomarkers for prognostics, treatment outcome prediction, and personalized medicine [28, 57, 44]. Additionally, GASTON introduces a new axis of tumor classification, in which tumors may be further characterized by the variation of distinct tumor processes across spatial gradients; for example, some tumors may have an increasing gradient of aerobic metabolism towards the tumor center (e.g. Figure 4) while other tumors may have a decreasing gradient. Another potential clinical implication is that the spatial gradients learned by GASTON could reveal spatial trajectories of *metastatic* migration, similar to how the spatial gradients learned by GASTON in the olfactory bulb show spatial trajectories of neural migration (Figure 5B). For example, the variation of EMT genes along the spatial gradients near the tumor boundary may reveal the molecular underpinnings of the *margination* process in which tumor cells migrate towards a vascular wall before metastasis [142, 165].

The inference of continuous variation in *transcriptomic* space, i.e. trajectory inference or *pseudotime* approaches, is widely applied in scRNA-seq analysis [49, 137, 107, 129]. Recently, there have been some attempts to adapt these approaches to SRT data [113, 51, 97, 85]. However, continuous variation in transcriptomic space is not equivalent to continuous variation in *physical* space that is modeled by isodepth. Indeed we find that existing approaches based on diffusion pseudotime [49] learn a coordinate that is nearly constant in each spatial domain, and thus obscures spatial variation in gene expression and cell type proportions within spatial domains (Figure S6). This limitation of existing scRNA-seq-based approaches underscores the need for methods like GASTON that model continuous *spatial* variation.

We note that the current derivation of isodepth by GASTON relies on two simplifying assumptions that may require adjustment for specific applications. First, we assume that all (spatially varying) genes share the same vector field of spatial gradients. Thus, GASTON will not automatically find multiple directions of spatial variation, where each direction corresponds to a subset of genes. For these situations, it might be appropriate to learn the isodepth using a restricted set of genes or a smaller region of a tissue slice; e.g. one may apply GASTON to spatial domains or gene sets obtained from a standard SRT or single-cell clustering algorithm. Second, we assume that the shared spatial gradient vector field is *conservative*, meaning that it does not “rotate” in space (i.e. curl (v) = 0). GASTON may not be applicable to tissue slices where this assumption is violated, although we are not aware of any such biological examples. An important next step would be to develop a framework for learning spatial gradients under relaxed mathematical assumptions, potentially using neural fields or transformers which have been used to learn vector fields in other areas of biology and machine learning [110, 152, 23].

We envision that the simplicity and generality of both the mathematical framework of the isodepth and the GASTON algorithm can be readily extended in several directions. First, the piecewise linear model of gene expression can be replaced by more complicated functions. While more complicated functions may be prone to overfitting with sparse SRT data, they may be appropriate for targeted SRT technologies —e.g. MERFISH [91], 10X Genomics Xenium [61], STARMap [143, 162], or NanoString CosMx [52] — that have higher detection efficiency. Second, it would be desirable to extend GASTON to identify 3-D spatial gradients, e.g. by utilizing spatial alignment tools [159, 77, 67, 64], as well as spatiotemporal gradients [114]. A third direction is to extend GASTON to other molecular modalities such as chromatin accessibility [164, 119] or protein/metabolite abundance [141, 82], e.g. using recent data on spatial measurements of ribosome-bound transcripts [161]. Fourth, there has been much work on quantifying transcriptomic vector fields by computing RNA velocity from ratios of spliced/unspliced RNA in single-cell RNA-seq data (e.g. [108, 70, 46, 11]) and it would be interesting to understand how RNA velocity varies along the spatial gradients learned by GASTON. Similarly, it would also be useful to understand how local microenvironments or cellular interactions, e.g. as learned by [150, 112, 30], vary along the GASTON spatial gradients. Finally, several recent papers have studied how genetic variants affect single-cell gene expression measurements, i.e. single-cell eQTLs (expression quantitative trait loci) [27, 95] and it would be useful to understand how genetic variants contribute to the continuous and discontinuous spatial gene expression patterns found by GASTON.

In summary, the topographic maps and gene expression functions computed by GASTON provide a novel and general framework for analyzing continuous and discontinuous spatial variation in gene expression from spatial sequencing data across many biological systems.

## 4 Methods

### 4.1 Modeling gene expression and spatial gradients

We derive a model for spatial domains and gradients of gene expression in spatially resolved transcriptomics (SRT) data. SRT technologies measure the expression of *G* genes in a tissue slice *T* ⊆ ℝ^2^, which we model with a *gene expression function* f : *T*→ℝ^*G*^. The vector f (*x, y*) = (*f*_1_ (*x, y*), …, *f*_*G*_ (*x, y*))^T^ gives the (normalized) expression of each gene at spatial location (*x, y*) in the tissue slice *T*, with the *g*-th component function *f*_*g*_ : ℝ^2^→ℝ describing the expression of a single gene *g*. For example, a gene *g* whose expression is constant across the tissue slice *T* has a constant expression function *f*_*g*_ (*x, y*) = *c*, while a gene that is differentially expressed in a region *R*⊆*T* might have the expression function *f*_*g*_ (*x, y*) = *c* ·1_ {(*x,y*) ∈ R}_ + *c*^′^·1 _ {(*x,y*) ∉*R*}._

We model each expression function *f*_*g*_ as a *piecewise continuous* function. Piecewise continuous functions model continuous spatial variation in gene expression while also allowing for discontinuities in expression due to sharp changes in cell type composition or other factors. We assume the expression functions *f*_*g*_ have the same pieces for all genes, and thus each expression function *f*_*g*_ has the form:

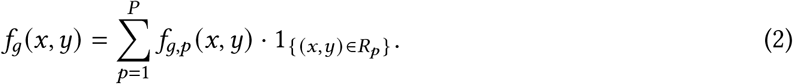

where *f*_*g,p*_ : ℝ^2^→ℝ are continuous functions and *R*_1_,…, *R*_*P*_⊆ℝ^2^ are a partition of the tissue slice *T* into *P* disjoint regions which we call *spatial domains*. Note that the spatial domains *R*_*p*_ need not be contiguous, and thus this model allows for physically separate locations within the tissue slice to contain a similar composition of cell types.

A *spatial gradient* describes how gene expression varies across the 2D tissue slice *T*. For a single gene *g*, the spatial gradient is given by the gradient ∇*f*_*g*_ of the expression function *f*_*g*_. More generally, the rows of the Jacobian matrix J(f) =[∇*f*_1_ … ∇*f*_*G*_]^T^ ∈ ℝ^*G*×2^ of the gene expression function f give the individual spatial gradient of each gene at each spatial location (*x, y*) ∈ *T*. Note that the rank of the Jacobian matrix J (f) is at most two.

Estimating the spatial gradients ∇*f*_*g*_ for every gene *g* from SRT data from a single tissue slice is difficult due to the limited spatial resolution and limited sequence coverage (e.g. sparsity) of the data. To avoid overfitting, we make some assumptions on the structure of the spatial gradients. Specifically, we assume that the Jacobian matrix **J**(**f**) has *rank one* at every spatial location (*x, y*) ∈ *T*, i.e. the rows *f*_*g*_ (*x, y*) of the Jacobian matrix **J**(**f**)(*x, y*) are linearly dependent for every spatial location (*x, y*)∈ *T*. This assumption is motivated by the observation that spatial expression gradients tend to be correlated; for example, many genes have been observed to have expression gradients along the same axes in the brain and liver [21, 9]. Under this assumption, for each spatial location (*x, y*) ∈ *T* there exists a vector v(*x, y*) ∈ ℝ^2^ such that the gradient vector ∇*f*_*g*_ (*x, y*) of each gene *g* is a scalar multiple of the vector v(*x, y*):

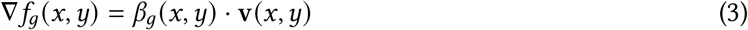

where β_*g*_ (*x, y*) : ℝ^2^→ℝ are scalar functions and v (*x, y*) is a vector field which we call the *spatial gradient vector field*. Since the expression function f is piecewise continuous, the gradient ∇*f*_*g*_ of each expression function *f*_*g*_ is also piecewise continuous, and so we may re-write (3) as

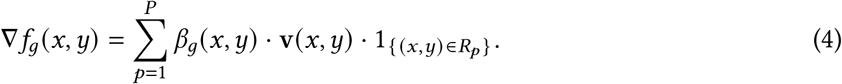

### 4.2 Conservative vector fields and piecewise linear functions

Equation (4) provides a general model for spatial gradients ∇*f*_*g*_ under a rank-one assumption on the Jacobian matrix **J (f)**. However, in practice, it is still difficult to estimate the parameters of (4) from SRT data, as we do not observe expression gradients ∇*f*_*g*_ but only the expression values *f*_*g*_. To derive a model for the expression functions *f*_*g*_ while minimizing overfitting, we make three simplifying assumptions on the spatial gradient vector field v, spatial domains *R*_*p*_, and scalar functions β_*g*_ (*x, y*).

First, we assume the spatial gradient vector field v is the gradient of a continuously differentiable, scalar function *d* : ℝ^2^→ℝ, i.e. v =∇*d*. We call *d* the *isodepth* of the tissue slice *T*. The isodepth *d* describes the “topography” of a tissue slice *T*, analogous to the elevation in a topographic maps of a geographic region. In physics, vector fields v that are the gradient of a scalar function *d* are called *conservative* vector fields, and the scalar function *d* is called the potential function as it measures potential energy at different locations in space, e.g. a gravitational potential function or an electric potential function [87]. In our setting, the scalar function *d* measures a *“gene expression potential”* at different locations in a tissue slice *T*. The vector field v being conservative is equivalent to the curl of v being 0 everywhere, i.e. there are no regions of the tissue where the vector field v “rotates”.

Second, we model each spatial domain *R*_*p*_ as a union of level sets of the isodepth *d*. Specifically, we assume that each spatial domain *R*_*p*_ ={(*x, y*): *b*_*p*−1_ *< d* (*x, y*) ≤*b*_*p*_} is equal to the set of spatial locations (*x, y*) with isodepth *d*(*x, y*) in the interval (*b*_*p*−1_, *b*_*p*_], for some real numbers−*∞* = *b*_0_ *< b*_1_ *<*…*< b*_*P*−1_ *< b*_*P*_ =*∞*. This ensures that the spatial domains *R*_*p*_ do not intersect, and leads to a particularly simple form for the expression function *f*_*g*_ as we show below.

Third, we assume that the scalar functions β_*g*_ (*x, y*) are constant inside each spatial domain *R*_*p*_; i.e., the scalar functions 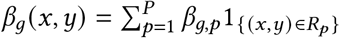 are piecewise constant.

Under these assumptions, the spatial gradients ∇*f*_*g*_ in (4) are equal to

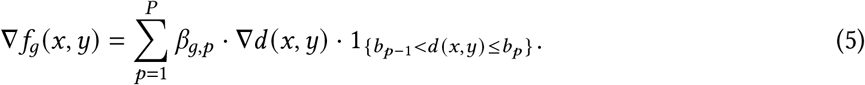

Integrating both sides of (5) yields the following closed form for the gene expression function *f*_*g*_:

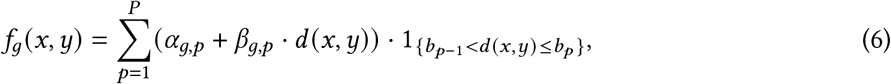

for some constants α_*g,p*_ and β_*g,p*_. Combining (6) for all genes *g* = 1,…, *G* yields the following expression for the gene expression vector **f** = (*f*_1_, …, *f*_*G*_):

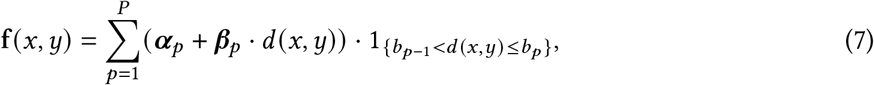

for vectors α_*p*_ = (α_*g,p*_)_*g*∈*G*_∈ ℝ^*G*^ and β_*p*_ = (β_*g,p*_)_*g*∈*G*_ ∈ ℝ^*G*^.

Thus, under our model, the gene expression function f (*x, y*) at spatial location (*x, y*) ∈ *T* is given by the composition f (*x, y*) = h(*d* (*x, y*)) of the isodepth *d* and a *piecewise linear* function h = (*h*_1_, …, *h*_*G*_) : ℝ → ℝ^*G*^ with *P* pieces and breakpoints *b*_1_, …, *b*_*P* −1_:

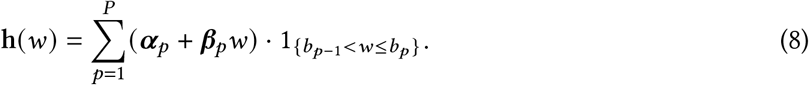

The vectors **α**_*p*_ and **β**_*p*_ are the *y*-intercepts and slopes, respectively, of the function h in the *p*-th piece across all *G* genes. We call the function h *w* a *one-dimensional (1-D)* expression function as it is a function of a single variable *w*, the isodepth, in contrast to the gene expression function f *x, y* which is a function of two spatial variables *x* and *y*.

#### Long-range spatial correlations and pooling

A major advantage of modeling gene expression as a function of isodepth is the ability to combine gene expression measurements from distinct spatial locations and thus overcome the sparsity of current SRT technologies. Specifically, all spatial locations with equal isodepth *d* have identical gene expression value h (*d)*, and so estimation of h (*d)* can use all locations on the contour of equal isodepth. This contour may traverse the entire tissue slice, and need not be a contiguous curve (e.g. Figure 2A). Thus, the isodepth model incorporates *“long-range”* spatial correlations [101], in contrast to many existing algorithms for analyzing SRT data which only incorporate local correlations between nearby spots, e.g. using hidden Markov random fields (HMRFs) [168, 34] or Gaussian processes (GPs) [135, 132, 130, 171]. Moreover, the isodepth model allows for “pooling” information across spatially separated regions of a tissue slice.

The isodepth model substantially generalizes the model of layered tissues and *“relative depth”* in [83] which restricted the spatial domains *R*_1_, …, *R*_*P*_ to be layers satisfying strict topological constraints. In contrast, here there are fewer topological constraints on the spatial domains *R*_1_, …, *R*_*P*_, and we learn the spatial domains and isodepth *de novo* from SRT data without any prior knowledge, as detailed below.

### 4.3 Maximum likelihood estimation

We compute the maximum likelihood estimators (MLEs) of the isodepth *d* and piecewise linear 1-D expression function h =(*h*_1_, …, *h*_*G*_) from SRT data. The observed SRT data consists of a transcript count matrix A = [*a*_*i,g*_] ∈ℝ^*N*×*G*^, where *a*_*i,g*_ is the UMI count of gene *g* in spot *i*, and a spatial location matrix S∈ℝ^*N*×2^, where each row s_*i*_ = (*x*_*i*_, *y*_*i*_)ℝ^2^ is the spatial location of the *i*-th spot. We define the Spatial Topography Problem (STP) as the following maximum likelihood estimation problem.

#### Spatial Topography Problem (STP)

*Given SRT data* (A, S) *and a number P of spatial domains, find a continuously differentiable function d* : ℝ^2^ → ℝ *and a piecewise linear function* h(*w*) : ℝ → ℝ^*G*^ *with P pieces that maximize the log-likelihood of the data:*

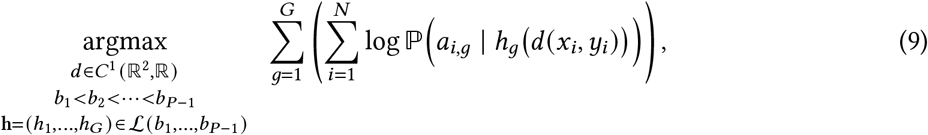

*where C*^1^ (ℝ^2^, ℝ) *is the space of continuously differentiable functions from* ℝ^2^ *to* ℝ *and* L(*b*_1_, …, *b*_*P* −1_) *is the set of piecewise linear functions with breakpoints b*_1_, …, *b*_*P* −1_.

The STP substantially generalizes the *L*-Layered Problem from our previous work [83], which assumed the isodepth *d* was given by a piecewise conformal map where the pieces are either bounded by lines or determined by prior knowledge on the shape of the spatial domains *R*_*p*_.

The STP is a challenging non-convex optimization problem over spaces of continuously differentiable and piecewise continuous functions. We solve this optimization problem using *deep learning*. By the universal approximation theorem [56], one can approximate a continuous function *d* with a neural network. Moreover, even a *piecewise* continuous function can be well-approximated by neural networks [79], although it may be computationally intractable to identify the individual pieces of the function [123]. Thus, we solve the STP in a two-step approach, where we first learn the isodepth *d* and then learn the piecewise linear expression function h.

#### Step 1

We estimate the isodepth *d* by solving a modified version of the maximum likelihood problem in (9) where we parametrize the functions *d* : ℝ^2^→ℝ and h = (*h*_1_, …, *h*_*G*_) : ℝ→ ℝ^*G*^ with neural networks with weights θ and θ ^′^, respectively.

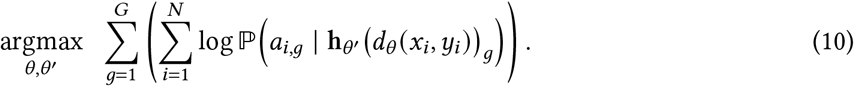

The modified problem in (10) is also a non-convex optimization problem for most neural network architectures. Nevertheless, by parametrizing the arguments with neural networks, we leverage the fact that such problems can be approximately and efficiently solved by modern deep learning frameworks such as PyTorch [102].

Solving (10) is equivalent to learning the parameters of a *single* neural network h_θ_′ ◦ *d*_θ_, where one of the hidden layers has only one hidden neuron whose value is the estimated isodepth *d*_θ_ (Figure 1). As a result, the isodepth corresponds to an *interpretable* hidden layer of a neural network.

Using the solution 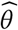 from (10) yields an estimate 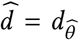 of the isodepth *d*. We expect the estimated isodepth *d* to be a good approximation of the solution to the STP (9), as both continuous functions *d* and piecewise continuous functions h can be well-approximated by neural networks [79]. However, it is difficult to identify the breakpoints *b*_1_, …, *b*_*P* −1_ — and thus the spatial domains *R*_*p*_ of the tissue slice — from the neural network h_θ_′. Therefore, we solve a second optimization problem to estimate the piecewise linear function h.

#### Step 2

We use the estimated 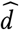 isodepth from Step 1 to estimate the piecewise linear function ĥ with breakpoints 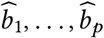 by solving the following optimization problem:

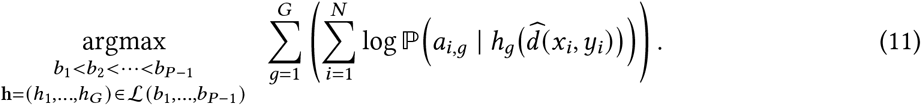

When there is only one gene, i.e. *G* = 1, then the maximum likelihood problem in (11) is an instance of segmented regression, a classical problem from statistics that is solved by dynamic programming (DP) [3, 7]. For *G >* 1 genes, we solve (11) using a variant of the segmented regression DP derived in [83].

### 4.4 Training and implementation

The algorithm described above can be implemented with different probability distributions 𝕡 (*a*_*i,g*_ | *f*_*g*_ (*x*_*i*_, *y*_*i*_)) = 𝕡 (*a*_*i,g*_ | *h*_*g*_ (*d* (*x*_*i*_, *y*_*i*_))) for the gene expression values *a*_*i,g*_. Following prior work [136, 120, 83, 100], we model the UMI counts *a*_*i,g*_ with a Poisson distribution of the form 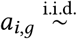 Pois (*U*_*i*_ · exp(*f*_*g*_ (*x*_*i*_, *y*_*i*_))), where *U*_*i*_ is the total UMI count at spot *i*. Another alternative is a Gaussian measurement model 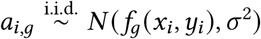.where σ^2^ is a shared variance parameter.

In practice, although one could use all or selected gene expression values instead, for efficiency we do not directly solve the STP (9) using the observed gene expression values but instead use the top generalized linear model principal components (GLM-PCs) [136]. This simplification is justified by our previous work [83] where we showed that for SRT data (A, S) generated from the Poisson expression model with a piecewise linear expression function h, then the top-2*P* GLM-PCs of the transcript count matrix A are also piecewise linear with Gaussian noise.

Specifically, we compute the top-2*P* GLM-PCs and solve (10) with these GLM-PCs under a Gaussian error model. For the colorectal tumor (Section 2.4), in order to capture spatial variation from the histological image, we use the top-(2*P*−3) GLM-PCs together with the mean R, G, and B values taken from the H&E stained image, resulting in (2*P*−3)+3 = 2*P* total features in the STP. We solve the optimization problem in (10) with neural networks *d*_θ_ and h_θ_′ that have two hidden layers of size 20 and are trained for 10000 epochs using the Adam optimizer [66]. Because of the non-convexity of (10), we use 30 random initializations and select the solution with the largest likelihood.

After solving (10) with the top-2 GLM-PCs and estimating the isodepth 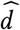, we then solve (11) with the top GLM-PCs to estimate the breakpoints 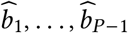. For most of the applications in this paper, we choose the number *P* of spatial domains using prior knowledge on the geometry of the tissue slice, e.g. for the cerebellum (Figure 2), we use *P* = 4 as prior work [19] showed that the cerebellum has four distinct layers. However, if the number *P* of domains is not known, then one may follow the model selection criteria used by [83], i.e. identifying an elbow in the log-likelihood plot, which we use for the DLPFC application (Figure S3).

Finally, we estimate the piecewise linear gene expression function ĥ by solving

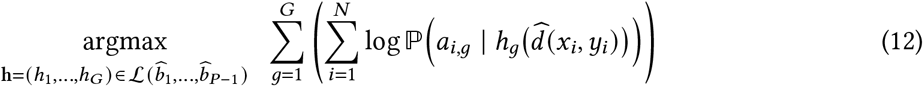

under the Poisson expression model for the UMI counts *a*_*i,g*_. We solve the optimization problem in (12) using Poisson regression with sklearn [103] for each individual gene *g* and spatial domain *R*_*p*_. To prevent overfitting, we subsequently perform a hypothesis test of whether each slope β_*g,p*_ of gene *g* in domain *R*_*p*_ is zero or non-zero, i.e. we test the hypotheses

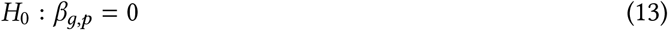

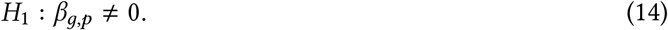

For each gene *g* and domain *R*_*p*_, we compute a log-likelihood ratio (LLR) for the null and alternative hypotheses under the Poisson expression model, and we estimate a *p*-value assuming that 2 · LLR follows a χ^2^-distribution, which holds asymptotically by Wilks’ theorem [147]. We set the slope β_*g,p*_ to zero if the *p*-value is less than 0.1.

Moreover, we estimate a 1-D expression function *h*_*g*_ only for genes *g* with at least *K* total UMI counts where *K* = 500 for the cerebellum and olfactory bulb (Sections 2.3, 2.5) and *K* = 1000 for the colorectal tumor (Section 2.4). These choices of *K* result in ≈2000−5000 genes for which we estimate an expression function. Moreover, for Slide-SeqV2 and Stereo-Seq applications with sparse UMI counts, we only estimate a slope β_*g,p*_ in domain *R*_*p*_ if there are at least *T* non-zero expression values in the domain. We use *T* = 75 for the cerebellum and *T* = 20 for the olfactory bulb, which are approximately 10% of the number of spatial locations in the smallest domain.

### 4.5 Quantifying spatial variation in gene expression

The piecewise linear expression functions 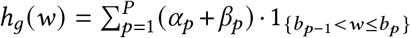reveal both discontinuities in expression and continuous variation within a domain, or intradomain variation, as we describe below.

#### Discontinuous expression

Let δ_*g,p*_ be the discontinuity of the function *h*_*g*_ at breakpoint *b*_*p*_, i.e. δ_*g,p*_ = (α_*g,p*+1_ + β_*g,p*+1_ · *b*_*p*_) − (α_*g,p*_ + β_*g,p*_ · *b*_*p*_). A large (absolute) discontinuity |δ_*g,p*_ | indicates a large discontinuous change in the expression of gene *g* at the boundary between spatial domains *R*_*p*_ and *R*_*p*+1_.

We say a gene *g* has a *discontinuity* in expression between spatial domains *R*_*p*_ and *R*_*p*+1_ if the esti mated discontinuity magnitude 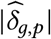 is greater than a threshold *t*_*p*_. We set the threshold *t*_*p*_ to be the tenth percentile of all estimated discontinuity magnitudes 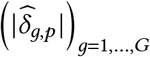 between spatial domains *R*_*p*_ and *R*_*p*+1_.

#### Intradomain variation

The slope β_*g,p*_ of the expression function *h*_*g*_ describes variation within a spatial domain *R*_*p*_. We say a gene *g* has *intradomain variation* in spatial domain *R*_*p*_ if the estimated magnitude 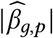 of the slope is greater than a threshold *s*_*p*_. That is, intradomain variation corresponds to a large *effect size* of the parameter β_*g,p*_. (Note that this effect size thresholding is distinct from the *p*-value thresholding in Section 4.4.) We set the threshold *s*_*p*_ to be the tenth percentile of all estimated slope magnitudes 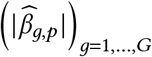 in domain *R*_*p*_.

### 4.6 Attributing continuous variation in expression to cell types

Intradomain variation in expression – i.e. a large magnitude of the slope β_*g,p*_ for a domain *R*_*p*_ in the piecewise linear fit – may be due to variation in expression within a cell type, variation in the proportions of cell types, or other biological causes. To illustrate, consider the 1-D expression function *h*(*w*) = *h*_*g*_ (*w*) for a single gene *g*. Given cell types *c* = 1, …, *C*, the function *h*(*w*) is given by

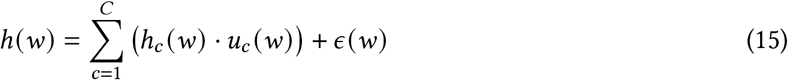

where *h*_*c*_ : ℝ→ ℝ is the *cell type c-specific* expression, 0≤*u*_*c*_ *w*≤1 is the proportion of cell type *c* at isodepth *w*, and ϵ *w* represents variation due to other factors.

Suppose that the expression function is *h*(*w*) = *e*· *u*_*c*_ (*w*) + ϵ (*w*); i.e. expression is constant for cell type *c* and zero for other cell types. If the cell type proportion *u*_*c*_ (*w*) or other variation function ϵ (*w*) are not constant functions of the isodepth (*w*), then the function *h w* will not be constant. Thus, when we fit the expression function *h* (*w*) with a piecewise linear function, we may estimate a non-zero slope β — reflecting variation in expression — even when there is no variation for any given cell type. This motivates the problem of learning *cell type-specific* expression functions *h*_*c,g*_ : ℝ→ ℝ for each gene *g* and cell type *c* which reveal whether variation is attributable to cell type or to other factors.

Here we derive a simple approach for estimating cell type-specific expression functions *h*_*c,g*_ from *single-cell resolution* SRT data with cell type annotations. Specifically, suppose we are given single-cell resolution SRT data (A, S) with cell type annotations *z*_*i,c*_ ∈ {0, 1}, where *z*_*i,c*_ = 1 if spot *i* contains cell type *c*, and *z*_*i,c*_ = 0 otherwise. We assume the isodepth *d* and breakpoints *b*_1_, …, *b*_*P* −1_ have already been computed as described in Section 4.4. We model the expression *a*_*i,g*_ at spatial location (*x*_*i*_, *y*_*i*_) with the Poisson expression model 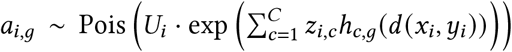 where *U*_*i*_ is the total UMI count at spatial location (*x*_*i*_, *y*_*i*_). Similar to Section 4.2, we model the cell type-specific expression functions h_*c*_ = (*h*_*c*,1_, …, *h*_*c,G*_) : ℝ → ℝ^*G*^ as piecewise linear functions of the form

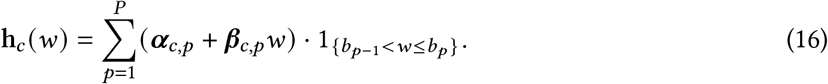

where α_*c,p*_ = (α_*c,p,g*_)_*g*=1,…,*G*_ and β_*c,p*_ = (β_*c,p,g*_)_*g*=1,…,*G*_ are the *cell type c-specific y*-intercepts and slopes, respectively, of the *cell type c-specific* expression function h_*c*_ = (*h*_*c*,1_, …, *h*_*c,G*_) in spatial domain *R*_*p*_.

The MLE of the piecewise linear, cell type *c*-specific expression functions h_*c*_ = (*h*_*c,g*_)_*g*∈*G*_ is given by

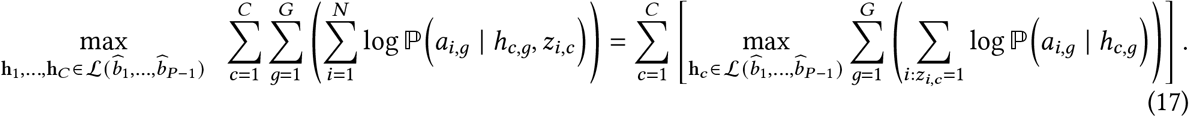

The inner optimization problem is an instance of the optimization problem in (12) restricted to spots *i* with cell type *c*, i.e. *z*_*i,c*_ = 1, and is solved using the same Poisson regression approach. Solving (17) yields estimated piecewise linear functions ĥ_*c,g*_ with *y*-intercept 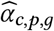 and slope 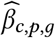 for each gene *g* in domain *R*_*p*_ and cell type *c*.

To assess whether intradomain variation is attributable to cell type, we compare the cell type *c*-specific slope β_*c,g,p*_ to the *cell type-agnostic* slope β_*g,p*_, which is derived from the *cell type-agnostic* expression function h *w* (Equation (8)). Specifically, we refer to the parameters α_*p*_ = (α_*g,p*) *g*∈*G*_ and β_*p*_ = (β_*g,p*) *g*∈*G*_ as the *cell type-agnostic y*-intercepts and slopes, respectively. If the cell type *c*-specific slope β_*c,g,p*_ is close or larger in magnitude to the *cell type-agnostic* slope β_*g,p*_, then the continuous variation in expression — i.e. the large value of β_*g,p*_ — is attributed to cell type *c*. Conversely, if the cell type-specific slope β_*c,g,p*_ is much smaller in magnitude than the cell type-agnostic slope β_*g,p*_, then the continuous variation in expression is not attributable to cell type *c*.

We quantify this intuition by dividing the genes with continuous variation identified in Section 4.5 into two groups based on the estimated cell type-specific slopes 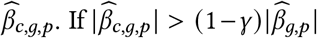 for some cell type *c* and a fixed constant γ, i.e. the magnitude of the cell type-specific slope 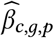 is close to or larger than the magnitude of the slope 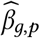, then we say that the expression variation within domain *R*_*p*_ is *attributable to cell type c*. On the other hand, if 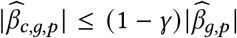 for all cell types *c*, then we say that there is *other* variation in the expression of gene *g* within domain *R*_*p*_. We use γ = 0.5 in our analyses.

### 4.7 Visualization

#### 4.7.1 Scaling isodepth to physical distance

The neural network in GASTON learns an isodepth *d* (*x, y*) that smoothly varies across a tissue slice *T*; however, the scaling of the learned isodepth *d* (*x, y*) is arbitrary. To improve the interpretability of the isodepth *d* (*x, y*) learned by the neural network, we scale the isodepth in each spatial domain to reflect approximate physical distances inside the domain. Briefly, we derive an estimate γ_*p*_ of the “average width” of each spatial domain *R*_*p*_ in μm, and we linearly transform the isodepth *d* (*x, y*) in each spatial domain such that the range of isodepth values in domain *R*_*P*_ is γ_*p*_.

We scale the isodepth in each spatial domain as follows. Given the isodepth *d* (*x, y*), spatial domains *R*_1_, …, *R*_*P*_, and breakpoints *b*_1_, …, *b*_*P* −1_ estimated from (10) and (11), we assume without loss of generality that the isodepth is linearly transformed such that min _*x,y* ∈*T*_ *d* (*x, y*) = 0 and max _*x,y* ∈*T*_ *d* (*x, y*) = 1, i.e. the breakpoints satisfy *b*_0_ = 0 *< b*_1_ *<*…*< b*_*P*−1_ *<* 1 = *b*_*P*_, where we set *b*_0_ = 0 and *b*_*P*_ = 1 for convenience. For each spatial domain *R*_*p*_, let γ_*p*_ be the average width of the domain, whose computation we describe below. We compute the *“scaled”* isodepth 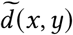 as

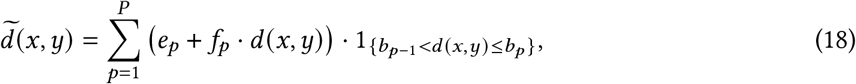

where *e*_*p*_, *f*_*p*_ are chosen such that 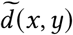 is continuous, and 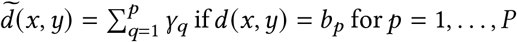. With this choice of *e*_*p*_, *f*_*p*_, the range of scaled isodepth values 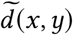 in a spatial domain *R*_*p*_ is given by

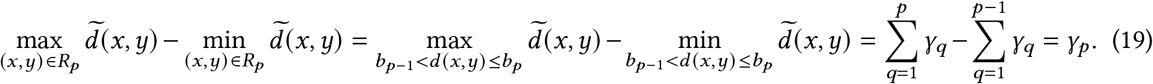

That is, the range of isodepth values 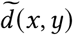 in each spatial domain is the average width γ_*p*_ of the domain *R*_*p*_.

We estimate the average widthγ_*p*_ of each spatial domain *R*_*p*_ by computing the median physical distance between the two boundaries of the domain *R*_*p*_. Specifically, let Γ_lower_ = {(*x*_*i*_, *y*_*i*_) ∈ *R*_*p*_ : *b*_*p*−1_ *< d* (*x*_*i*_, *y*_*i*_) *< b*_*p*−1_+ϵ and let Γ_upper_ = {(*x*_*i*_, *y*_*i*_) ∈ *R*_*p*_ : *b*_*p*_−ϵ^′^*< d* (*x*_*i*_, *y*_*i*_)*< b*_*p*_} be the set of spatial locations on the lower and upper boundary curves of the spatial domain *R*_*p*_, respectively. We set γ_*p*_ to be the median distance between ssseach spot (*x, y*) ∈Γ_lower_ and the closest spot in Γ_upper_ We choose ϵ, ϵ^′^ such that Γ_lower_ and Γ_upper_ visually correspond to the spatial domain boundaries.

For 10x Genomics Visium data, we multiply each average width γ_*p*_ by 100, since the physical distance between the centers of adjacent spots in the 10x Visium slide is 100μm. For Slide-seqV2 data, we multiply each average width γ_*p*_ by 64/100, since two beads that are 100 pixels apart in the Slide-SeqV2 microscopy image have a physical distance of roughly 64μm [116].

#### 4.7.2 Visualizing 1-D expression functions

To simplify the visualization of the 1-D expression functions h, we aggregate the counts *a*_*i,g*_ for spots (*x*_*i*_, *y*_*i*_) with approximately equal isodepth values *d* (*x*_*i*_, *y*_*i*_), as in [83]. Specifically, we partition the range of isodepth values into a union *B*_1_ ∪ … ∪ *B*_*M*_ of intervals *B* _*j*_, and we compute the total expression value 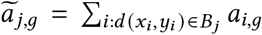 for gene *g* in each interval *B* _*j*_. We call ã _*j,g*_ the *pooled* expression value of gene *g* at pooled spot *j*. Pooling does not affect inference of the 1-D expression function h in the STP, as the function h obtained by maximizing the log-likelihood (9) with pooled data is equal to the function obtained by maximizing (9) with the original data, as shown in [83].

We plot expression as log pooled counts per million (CPM) 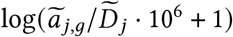, where 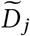 is the sum of the total UMI counts across all spots in the *j* th pooled spot. The log pooled CPM has approximately the same scale as the expression function *h*_*g*_ (*w*) + log(10^6^) for each gene *g*.

### 4.8 Marker gene analysis

For the marker gene comparison in Section 2.3, we derived a ranking of domain specific marker genes from the GASTON inferred 1-D expression functions *h*_*g*_ by ranking genes by the standard deviation of the mean of each expression function. Specifically, for each gene *g*, we compute the mean *m*_*g,p*_ of the 1-D expression function *h*_*g*_ (*w*) in spatial domain *R*_*p*_, i.e. 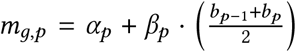, and we rank genes by the standard deviation of the values *m*(_*g,p*) *p*=1,…,*P*_. Intuitively, a marker gene should have high expression in one spatial domain and low expression in other domains, leading to a large standard deviation, while a non-marker gene will have similar expression in all domains, leading to a small standard deviation.

### 4.9 Spatial coherence score

We quantify the spatial coherence of domain labels using a score based on O’Neill’s spatial entropy measure [98, 4] which has previously been used to quantify spatial coherence in SRT data [159]. The spatial entropy measures the fraction of neighboring spots having the same label compared to random assignments of labels. A large spatial entropy indicates that the distribution of labels of neighboring spots is close to the uniform distribution, i.e the labels are spatially coherent, whereas a small spatial entropy indicates that nearby spots frequently have the same label, i.e. the labels are spatially coherent.

We use a modified version of the spatial coherence score used by [159] that is scaled to lie in [0, 1]. Specifically, following the notation in [159], we define the spatial coherence score as 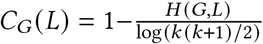.

### 4.10 Data collection and method details

For the cerebellum analysis in Sections 2.2 and 2.3, we used replicate 1 from the RCTD/C-SIDE data repository [18]. Figure 2J was created with BioRender.com. For the marker gene comparison in Figure 3A, we derived a gene ranking for each method and evaluated the AUPRC compared to known marker genes of the oligodendrocyte, granule, Purkinje, Bergmann, and molecular cell types in the cerebellum. These marker genes were the combination of cell type marker genes from PanglaoDB [40], the Allen Mouse Brain Atlas [71], Harmonizome [118], and the supplement of [69].

We obtained the olfactory bulb SRT dataset from [42]. We obtained cell type annotations for each spot in the tissue (Figure 5D) by using scANVI [154] to integrate the SRT data with a separate mouse olfactory bulb scRNA-seq dataset [133]; for the scRNA-seq data, we followed the pre-processing steps in [76].

#### SpiceMix

We followed the Visium Jupyter notebook tutorial on Github with parameters *K* = 6 (for the *K*-NN graph) and n neighbors=200.

#### Non-negative spatial factorization (NSF)

We followed the Github tutorial and trained for 150 iterations to obtain 10 factors. Since NSF identifies factors rather than spatial domains, we identified NSF spatial domains by using the NSF factors as input for the Louvain clustering module from SpiceMix [24].

#### RCTD/C-SIDE

For the cerebellum analysis, we used the cell type labels provided in the RCTD data repository. We followed the C-SIDE tutorial to identify cell type-specific differentially expressed genes. We ran two versions of C-SIDE: (1) without any covariates, and (2) with the isodepth *d* (*x, y*) as a covariate for each spatial location (*x, y*). For the analysis in Section 2.3, we ranked genes by their minimum C-SIDE *p*-value across all cell types.

#### SpaGCN

We ran SpaGCN following the Github tutorial. For the analysis in Section 2.3, we used a ranking where the SpaGCN spatially varying genes are tied for first and all other genes are tied for second.

#### HotSpot

We ran HotSpot following the tutorial here. For the analysis in Section 2.3, we ranked genes according to their *p*-value.

#### trendsceek*

We used the Seurat implementation of trendsceek as described here. For the analysis in Section 2.3, we ranked genes according to their *p*-value.

#### SpatialDE

We ran SpatialDE following the Github example. For the analysis in Section 2.3 we ranked genes according to their *p*-value.

#### SPARK-X

We ran Spark-X following the tutorial here. For the analysis in Section 2.3 we ranked genes according to their *p*-value.

#### SpaceFold

We ran SpaceFold following the Github example code.

## Supporting information

GASTON piecewise linear fits for cerebellum sample

GASTON piecewise linear fits for olfactory bulb sample

GASTON piecewise linear fits for colorectal tumor sample

## Acknowledgements

This research is supported by National Cancer Institute (NCI) grants U24CA248453 and U24CA264027 to B.J.R. U.C. was supported by a National Science Foundation Graduate Research Fellowship and the Siebel Scholars program. B.J.A. gratefully acknowledges financial support from the Schmidt DataX Fund at Princeton University, made possible through a major gift from the Schmidt Futures Foundation. H.S. is funded by the Princeton Ludwig Branch. C.M. is a Damon Runyon Fellow supported by the Damon Runyon Cancer Research Foundation (DRQ-15-22).

## Data and code availability

This paper analyzes existing, publicly available data. The cerebellum SRT dataset was obtained from [18]; the olfactory bulb SRT data set was obtained from [42]; the colorectal tumor SRT dataset was obtained from [149]; and the DLPFC SRT dataset was obtained from [89]. The code for GASTON is publicly available at https://github.com/raphael-group/GASTON.

## Supplemental Information

### A Expression models and pooling

We assume the UMI counts *a*_*i,g*_ follow the Poisson expression model, i.e. the UMI counts *a*_*i,g*_ are independent and follow a Poisson distribution of the form 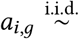 Pois (*U*_*i*_ · exp (*f*_*g*_ (*x*_*i*_, *y*_*i*_))) where *U*_*i*_ is the total UMI count at spot *i*.

Suppose the isodepth *d* is known, and let γ_1_, …, γ_*N*_ ′ be the unique isodepth values *d* (*x*_*i*_, *y*_*i*_) across all spots s_*i*_ = (*x*_*i*_, *y*_*i*_). Let *B* _*j*_ = {*i* : *d* (*x*_*i*_, *y*_*i*_) = γ _*j*_} be the set of spots with isodepth equal to γ _*j*_. Let 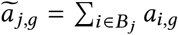 be the total expression for gene *g* over all spots *i* ∈ *B* _*j*_, i.e. ã_*j,g*_ is the total expression for all spots with isodepth γ _*j*_. We say *B* _*j*_ is a *pooled* spot and we call ã_*j,g*_ the *pooled* expression of gene *g* at the *j* -th pooled spot.

The solution to the MLE problem in (9) with isodepth *d* is equal to the solution of the following optimization problem

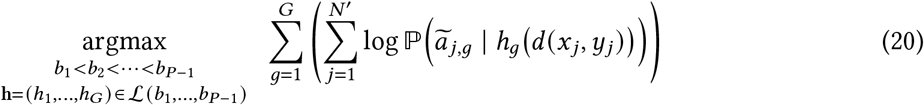

where the inference is performed with pooled expression values ã _*j,g*_. Thus, one obtains the same expression function h whether one computes the MLE (9) over all data points, or first sums spots with the same isodepth, i.e. *pooling* spots by their isodepth, and then computes the MLE. See [83] for more details.

### B Dimensionality reduction using GLM-PCA

Given SRT data A, S, we first run GLM-PCA (generalized linear model principal components analysis) [136] and obtain the top-2*P* GLM-PCs u_*j*_ = [*u*_*i,j*_] ∈ ℝ^*N*^ for *j* = 1, …, 2*P*. Next, we compute the MLE in (9) using these PCs and a Gaussian error model, i.e. we solve

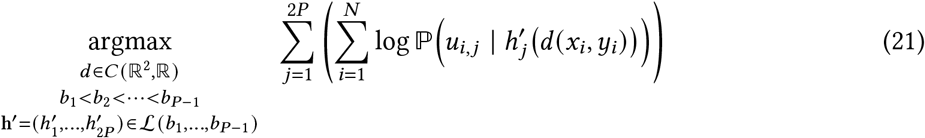

with 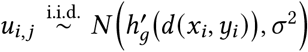 for some shared variance parameter σ^2^. (Note that the value of the variance σ^2^ does not affect the solution to (21).) Solving (21) an estimated isodepth 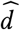 and breakpoints 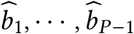.Finally, we solve the MLE problem in (9) fixing the estimated isodepth 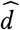 and breakpoints 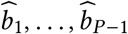 i.e.

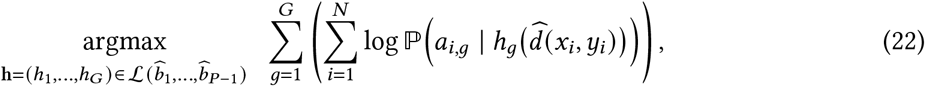

where we assume the UMI counts *a*_*i,g*_ follow the Poisson expression model described above. Solving (22) is equivalent to solving *G* · *P* Poisson regression problems, one problem for each combination of the *G* genes and *P* spatial clusters.

## C Comparison to SpaceFlow

SpaceFlow [113] learns a 1-D coordinate, which they call a *pseudo-Spatialtemporal Map (pSM)*, at each spatial location in a tissue by running diffusion pseudotime [49] on embeddings obtained from a graph neural network. We compared the SpaceFlow pSM to the GASTON isodepth on the mouse cerebellum SRT data from Section 2.2 (Figure S1A,B) and the mouse olfactory bulb SRT data from Section 2.5 (Figure S6C,D). Visually, the isodepth learned by GASTON varies continuously in the tissue while the SpaceFlow pSM does not. For example, in the cerebellum, the pSM is constant — and thus does not continuously vary — within each spatial domain, e.g. in the granule layer, the contours of the isodepth (Figure S1C) smoothly vary while the contours of the pSM (Figure S1D) are irregular. In the olfactory bulb, the pSM is constant in the interior of the tissue (Figure S6C).

We quantify the continuous variation within each layer using the quartile coefficient of dispersion (QCOD) [15], a robust statistic measuring the variation of a dataset, with a large QCOD indicating a larger degree of variation in the data. We first scale the isodepth and the pSM to be in [0, 1] so that they have the same measurement scale; moreover, before computing the QCOD within each layer, we shift the measurements to have the same mean in order to guarantee that the QCOD values are comparable. We observe that in the cerebellum, GASTON has larger QCOD than SpaceFlow in three out of four spatial domains (Figure S1E), indicating that there is substantially more spatial variation in the GASTON isodepth compared to the SpaceFlow pSM. Similarly, in the olfactory bulb, GASTON has larger QCOD than SpaceFlow in six out of seven domains (Figure S6B).

## D DLPFC comparison

We evaluated GASTON on SRT data from the human dorsolateral prefrontal cortex (DLPFC) measured with 10x Visium [89]. We analyzed eight DLPFC tissue slices from two donors. These slices were manually annotated with the six layers of the DLPFC and white matter (WM) and have a curved, layered geometry, providing spatial structure that may help GASTON accurately learn the geometry of these tissue slices. We compared the spatial domains identified by GASTON to two graph deep learning approaches, SpaGCN [58] and STAGATE, and our previous method Belayer [83], which requires supervision in the form of approximate layer boundaries. We evaluated each method by computing the adjusted Rand index (ARI) between the estimated spatial domains and the manually annotated layers.

GASTON achieves a higher average AUPRC than the graph deep learning methods SpaGCN and STA-GATE (Figure S3A). Moreover, despite being completely unsupervised, GASTON has comparable AUPRC to Belayer, which requires supervision (Figure S3A,B). Importantly, the isodepth*d* (*x, y*) learned by GASTON (Figure S3C) is highly correlated with the “relative depth” 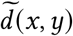 that Belayer estimates by solving the heat equation with known layer boundaries (Figure S3D), demonstrating that the neural network used by GASTON indeed learns the cortical depth of each layer. On the other hand, the isodepth *d* has lower correlation (Figure S3D) with both the top principal component (PC1) and the top generalized linear model principal component (GLM-PC1), which are derived solely from gene expression and do not use the spatial coordinates. These comparisons indicate the importance of spatial information in deriving an accurate measurement of layer depth.

Overall, the improved performance of GASTON demonstrates the value of using simple and interpretable neural network architectures.

**Figure S1:**
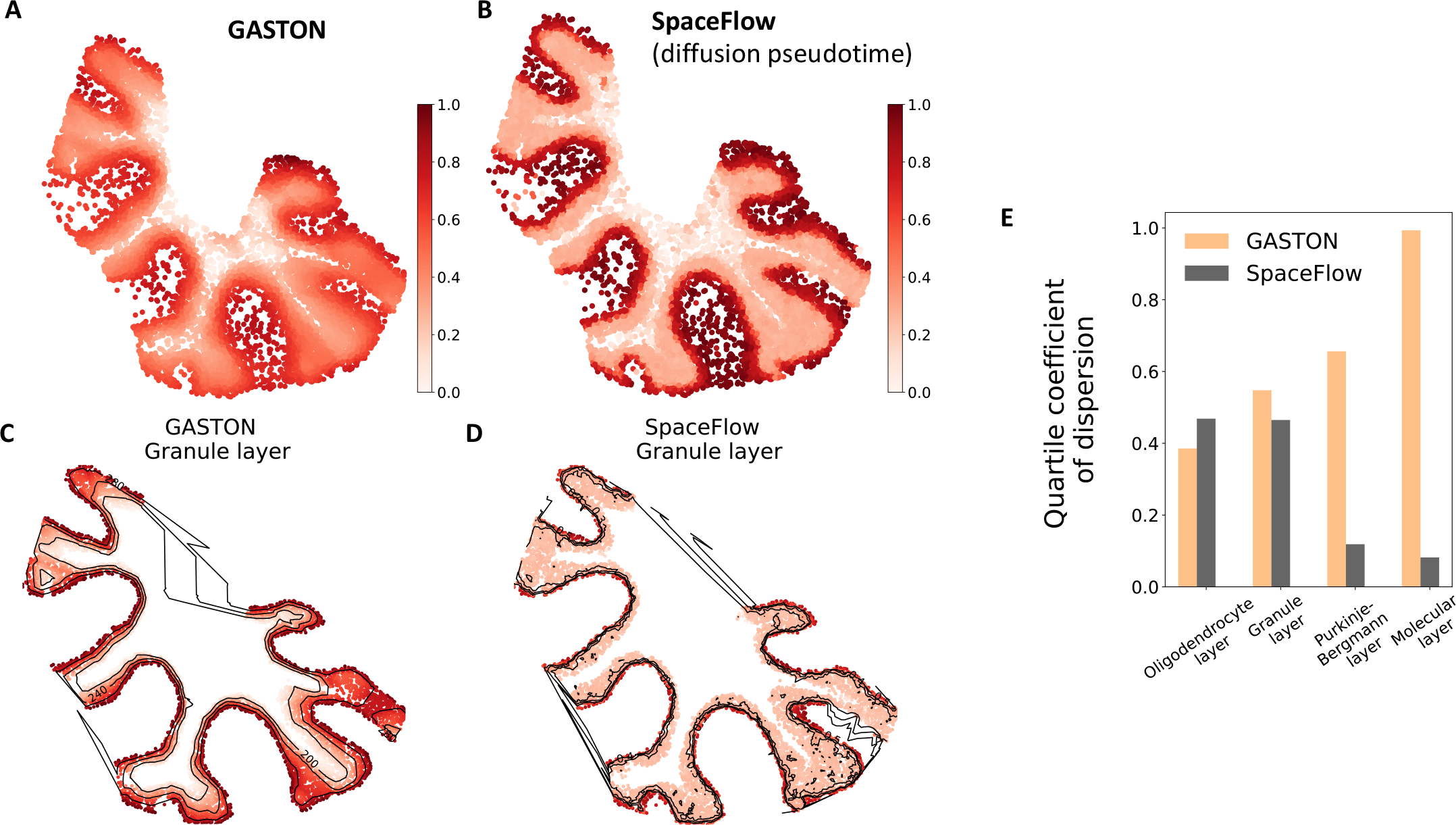
**(A)** The isodepth *d* (*x, y*) learned by GASTON scaled to [0, 1]. **(B)** The pseudo-Spatiotemporal Map (pSM) learned by SpaceFlow [113] scaled to [0, 1]. **(C)** The isodepth *d* (*x, y*) in the granule layer (as identified by GASTON), shown with three equally spaced contours of equal isodepth. **(D)** The pSM in the granule layer shown with three equally spaced contours of equal pSM. **(E)** The quartile coefficient of dispersion of the GASTON isodepth and the SpaceFlow pSM in each layer of the cerebellum.

**Figure S2:**
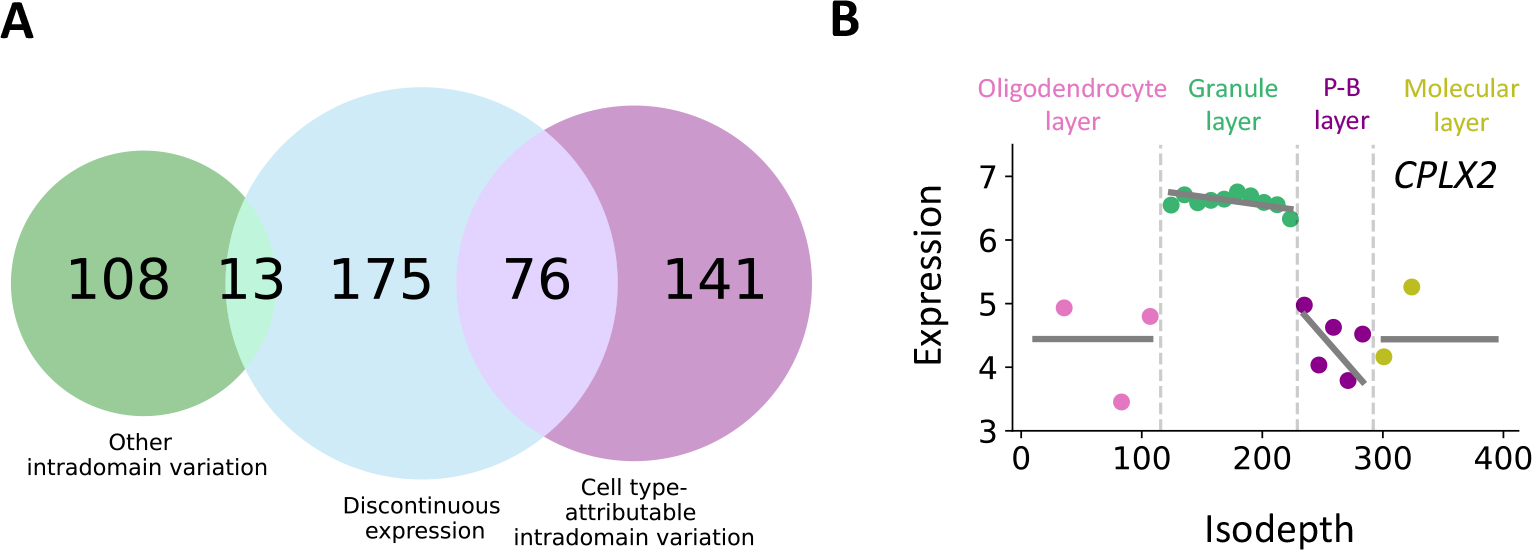
**(A)** Venn diagram of spatially varying genes identified by GASTON in the mouse cerebellum. Numbers indicate genes with specified spatial expression pattern(s). **(B)** Isodepth versus expression for *CPLX2*, which has discontinuities in expression at the granule layer boundaries.

**Figure S3:**
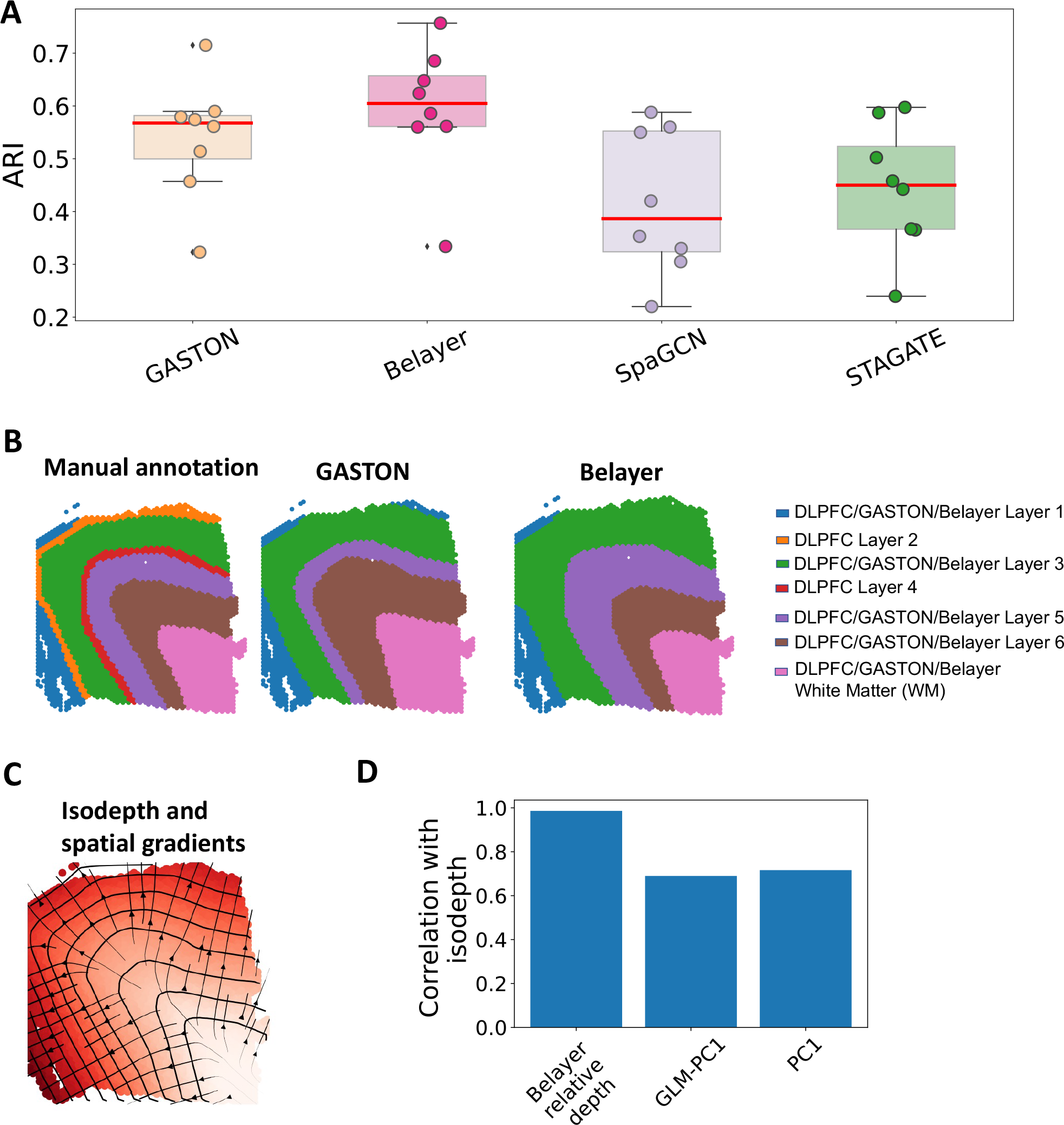
**(A)** Adjusted rand index (ARI) for GASTON, Belayer [83], SpaGCN [58], and STAGATE [32] in identifying the spatial domains of the dorsolateral prefrontal cortex (DLPFC). **(B)** The manually annotated domains and the domains identified by GASTON and Belayer for DLPFC sample 151673. **(C)** Isodepth *d* and spatial gradients ∇*d* learned by GASTON for DLPFC sample 151673. **(D)** Correlation between the GASTON isodepth and (1) the relative depth 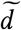estimated by Belayer using prior knowledge of the layer boundaries (Belayer relative depth); (2) the first generalized linear model principal component (GLM-PC1); and (3) the first principal component (PC1).

**Figure S4:**
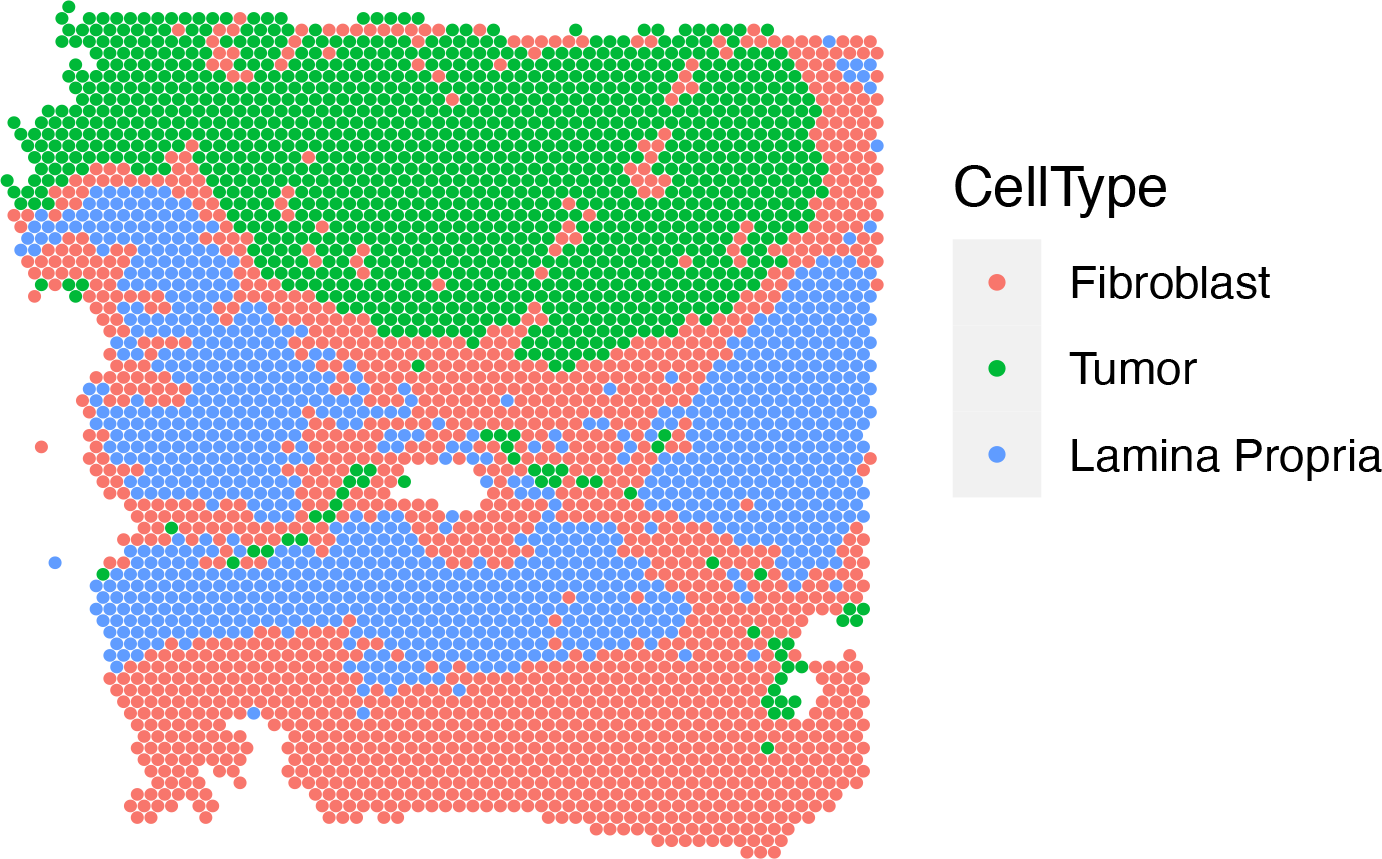
Cell type labels for each spot in 10x Genomics Visium data from a colorectal tumor slice derived in the original study [149] using Seurat [17].

**Figure S5:**
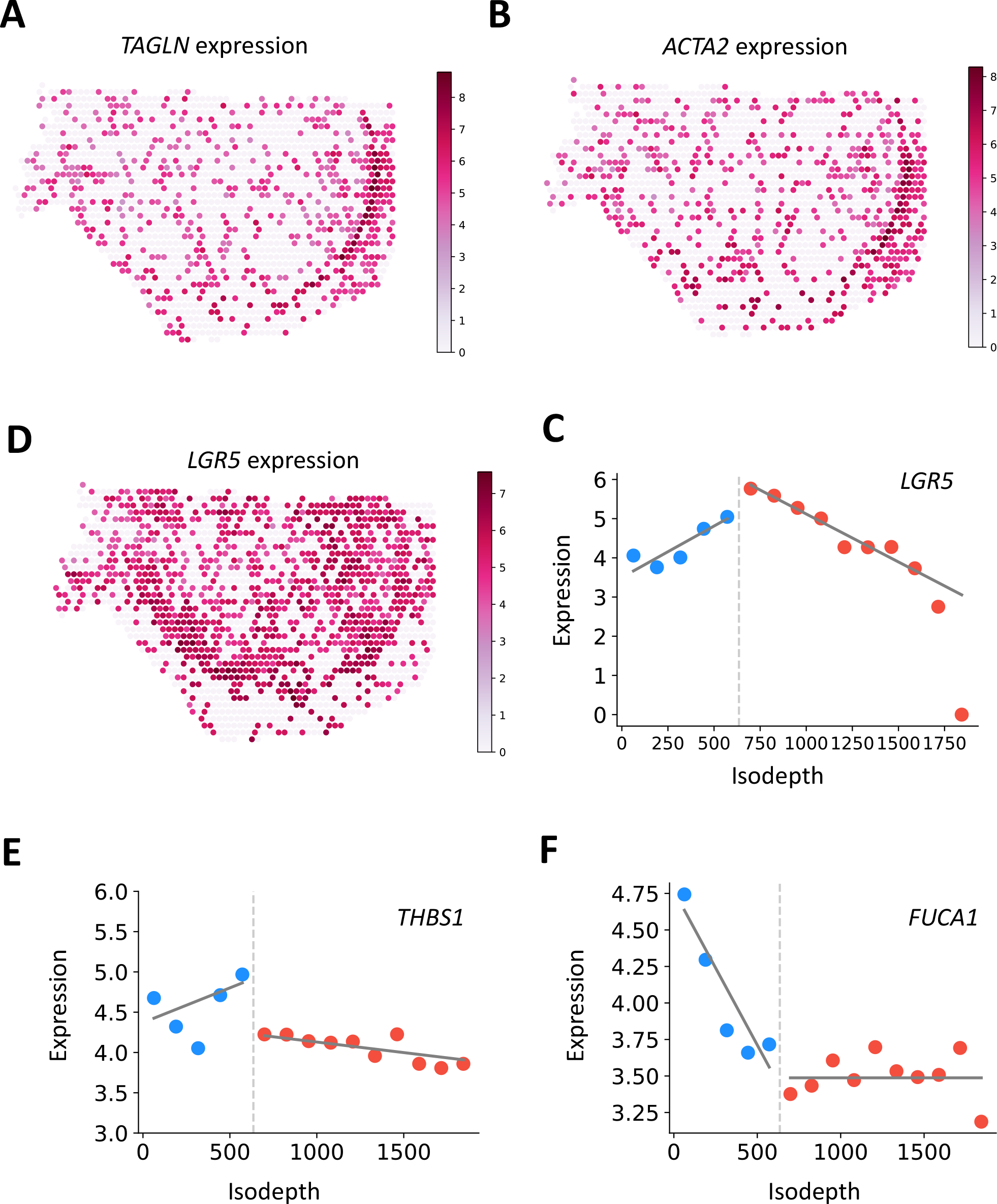
**(A-C)** Expression shown in log CPM for Type II genes **(A)** *TAGLN*, **(B)** *ACTA2*, and **(C)** *LGR5*. **(D-F)** Expression versus isodepth for Type II gene **(D)** *LGR5* and Type III genes **(E)** *THBS1* and **(F)** *FUCA1*.

**Figure S6:**
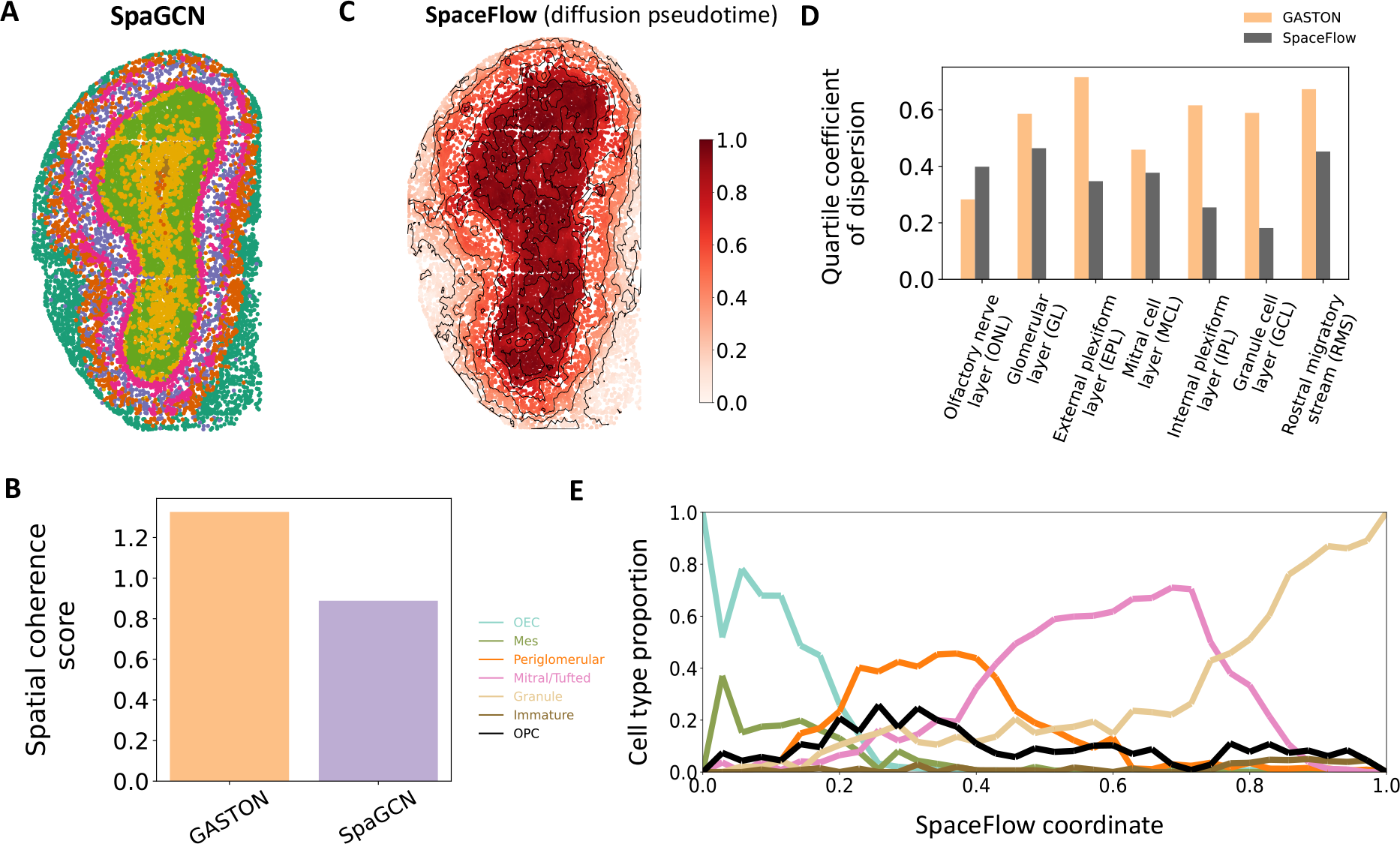
**(A)** Spatial domains learned by SpaGCN [58]. **(B)** Spatial coherence of spatial domains identified by GASTON (Figure 5C) and SpaGCN. **(C)** Pseudospatial-temporal map (pSM) learned by SpaceFlow [113], which utilizes the scRNA-seq based method diffusion pseudotime [49]. Curves denote contour lines of equal pSM. **(D)** Quartile coefficient of dispersion of the GASTON isodepth and the SpaceFlow pSM in each spatial domain identified by GASTON. **(E)** Cell type proportion as a function of SpaceFlow pSM.

**Figure S7:**
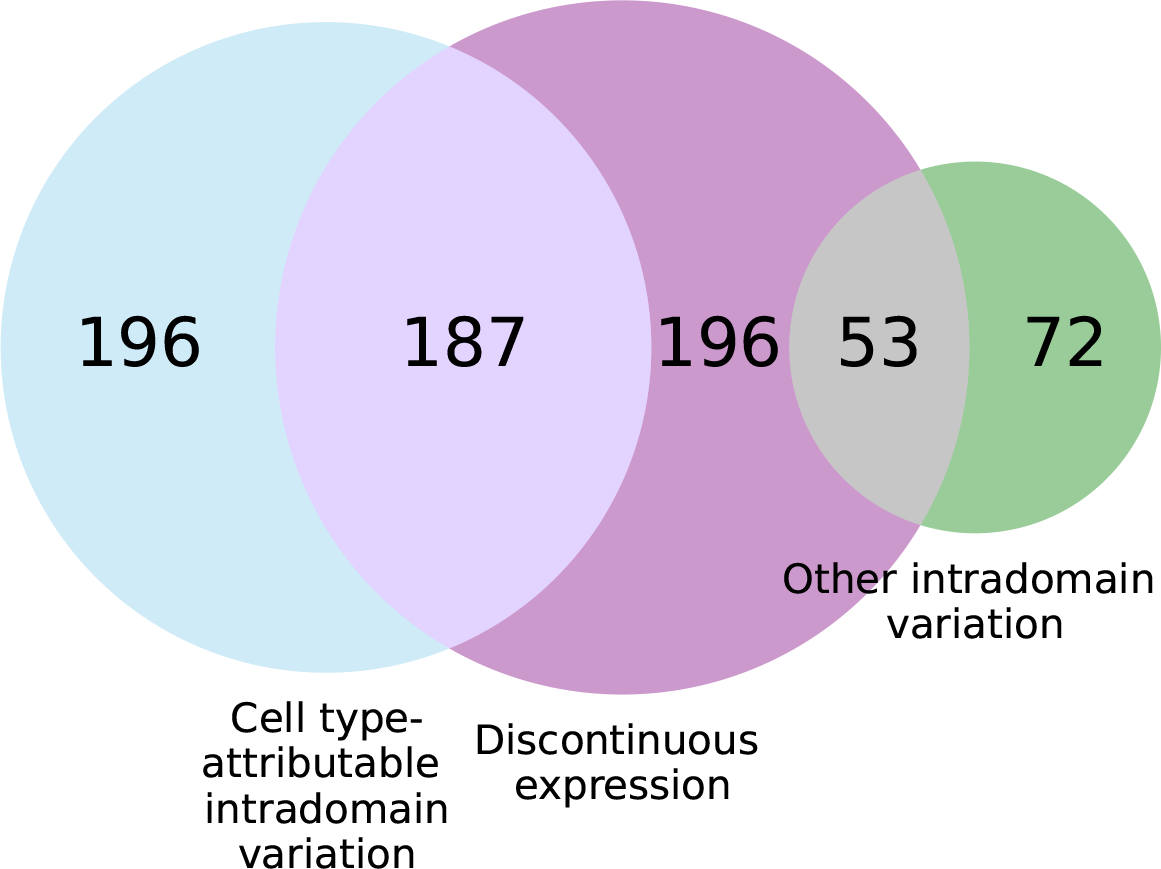
Venn diagram of spatially varying genes identified by GASTON in the olfactory bulb.

